# Mice With Monoallelic *GNAO1* Loss Exhibit Reduced Inhibitory Synaptic Input To Cerebellar Purkinje Cells

**DOI:** 10.1101/2021.09.23.461583

**Authors:** Huijie Feng, Yukun Yuan, Michael R. Williams, Alex J. Roy, Jeffery Leipprandt, Richard R. Neubig

## Abstract

*GNAO1* encodes Gα_o_, a heterotrimeric G protein alpha subunit in the G_i/o_ family. In this report, we used a *Gnao1* mouse model “G203R” previously described as a “gain-of-function” *Gnao1* mutant with movement abnormalities and enhanced seizure susceptibility. Here, we report an unexpected second mutation resulting in a loss-of-function Gα_o_ protein and describe alterations in central synaptic transmission.

Whole cell patch clamp recordings from Purkinje cells (PCs) in acute cerebellar slices from *Gnao1* mutant mice showed significantly lower frequencies of spontaneous and miniature inhibitory postsynaptic currents (sIPSCs and mIPSCs) compared to WT mice. There was no significant change in sEPSCs or mEPSCs. Whereas mIPSC frequency was reduced, mIPSC amplitudes were not affected, suggesting a presynaptic mechanism of action. A modest decrease in the number of molecular layer interneurons was insufficient to explain the magnitude of IPSC suppression. Paradoxically, G_i/o_ inhibitors (pertussis toxin), enhanced the mutant-suppressed mIPSC frequency and eliminated the difference between WT and *Gnao1* mice. While GABA_B_ receptor regulates mIPSCs, neither agonists nor antagonists of this receptor altered function in the mutant mouse PCs. This study is the first electrophysiological investigation of the role of G_i/o_ protein in cerebellar synaptic transmission using an animal model with a loss-of-function G_i/o_ protein.

**Significance Statement:** This is the first report on the electrophysiological mechanisms of a movement disorder animal model with monoallelic *Gnao1* loss. This study illustrates the role of Gα_o_ protein in regulating GABA release in mouse cerebellum. This study could also facilitate the discovery of new drugs or drug repurposing for *GNAO1*-associated disorders. Moreover, since *GNAO1* shares pathways with other genes related to movement disorders, developing drugs for the treatment of *GNAO1*-associated movement disorders could further the pharmacological intervention for other monogenic movement disorders.

## Introduction

*GNAO1* encodes the α subunit of a heterotrimeric G protein, G_o_, the most abundant membrane protein in the brain. It participates in multiple neural signaling pathways (Jiang & Bajpayee, 2009). The G_o_ protein functions in a broad range of signaling pathways including inhibition of cAMP production (Sunahara & Taussig, 2002), Gβγ-mediated inhibition of synaptic vesicle release (Zurawski et al., 2017), suppression of N- and P/Q-type calcium channel-mediated currents (Ikeda, 1996), and stimulation of G-protein regulated inward rectifying potassium (GIRK) channels (Zhang, Dickson, & Doupnik, 2004). Mutations in *GNAO1* were found in early onset epileptic encephalopathy (DEE17; OMIM#615473; Nakamura et al., 2013) and neurodevelopmental disorder with involuntary movements (NEDIM; OMIM#617493), a rare neurogenetic disorder, characterized by early onset of hypotonia, movement disorder and developmental delay. A genotype-phenotype correlation of *GNAO1*-associated neurological disorders, based on G_o_’s inhibition of cAMP production, showed that loss-of-function (LOF) and partial-loss-of-function (PLOF) *GNAO1* mutations are mainly associated with epilepsy, while the gain-of-function (GOF) and normal-function (NF) mutations are generally found in movement disorder patients (Feng et al., 2017). A mouse model with a knock-in GOF mutation G203R was reported (Feng et al., 2019). However, based on new genotyping data for the *Gnao1* G203R mutant mice, we have discovered a second mutation within the *Gnao1* gene. The second mutation disrupts a splice site and likely results in a haploinsufficient LOF allele for *Gnao1*. These mice were previously shown to exhibit movement abnormalities, assessed in behavioral tests including RotaRod, grip strength, and DigiGait, and showed increased sensitivity to pentylenetetrazol (PTZ) kindling (Feng et al., 2019).

To examine the possible mechanisms of movement abnormalities of the *Gnao1* mutant mice, we examined changes in spontaneous synaptic responses in Purkinje cells (PCs) in acute cerebellar slices isolated from WT and *Gnao1* mutant mice using whole-cell patch clamp recording techniques.

The cerebellum has long been known for its critical regulation of motor coordination; its role in ataxia has been well-studied (Buijsen, Toonen, Gardiner, & van Roon-Mom, 2019). Recent findings show that structural and/or functional abnormalities of the cerebellum are also associated with dystonia (Bologna & Berardelli, 2018) and chorea (Walker, 2016). Several reports demonstrated a role of the cerebellum in DYT1 hereditary dystonia (Fremont, Tewari, Angueyra, & Khodakhah, 2017; Vanni et al., 2015). Interestingly, dystonia, chorea/athetosis and ataxia are common movements abnormalities in *GNAO1* patients (Arya, Spaeth, Gilbert, Leach, & Holland, 2017; Kelly et al., 2019; Schorling et al., 2017). Structurally, a core circuit in the cerebellum mediates all its functions (Eccles, 1967; Reeber, Otis, & Sillitoe, 2013). This circuit centers on PCs, the axons of which are the sole output of the cerebellar cortex (Brown et al., 2019). Each PC receives two major glutamatergic excitatory inputs: climbing fibers (CFs) and parallel fibers (PFs), and two sets of GABAergic inhibitory inputs from molecular layer internuerons (stellate and basket cells). We thus investigated whether GNAO mutant mice demonstrated alteres synaptic input to PCs.

We report here decreased inhibitory input from cerebellar presynaptic GABAergic terminals upon PCs; there was a reduced frequency of sIPSCs and especially mIPSCs in the mutant mouse cerebellar slices but no change in EPSCs. A likely mechanism underlying this reduction in spontaneous release of GABA is that a LOF G_o_ mutant frees up more Gβγ to mediate presynaptic inhibition. Interestingly, the GABA_B_ receptor, which selectively controls mIPSCs, does not appear to be involved in the action of the mutant protein. There were no significant changes in inhibitory synapse morphology upon PCs and a modest decrease in molecular layer interneuron count which is unlikely to solely account for the reduction of IPSCs. These observations, and the reduced Gα_o_ expression in brain, with the normal levels of Gβγ. strongly support a model with a haploinsufficient LOF *Gnao1* mutant which mainly functions through the receptor-independent release of free Gβγ subunits in neurons to suppress the activity of inhibitory neurons in the cerebellum of the *Gnao1* mutant mice.

## Materials and Methods

All animal procedures complied with the U.S. National Institutes of Health guidelines on animal care and were approved by the MSU Institutional Committee on Animal Care and Use. The G203R mutation was introduced into the *Gnao1* gene of the C57BL/6J mice by CRISPR-Cas9 and was previously considered to be a gain-of-function (GOF) mutation (Feng et al., 2019). Here, we identity an unexpected mutation that changes two bases 13 and 14 bp upstream of the designed G203R mutation (Figure 1). This converts the AGG at the splice acceptor site on exon 6 to an AAA which does not splice correctly. RT-PCR analysis and sequencing showed that there was virtually not correctly spliced mutant *Gnao1* RNA that contained the G203R mutant allele. Western blots showed that the mutant mice had reduced expression of Gα_o_ protein thus resulting in a haploinsufficient LOF mechanism. Thus, in this report, we refer these mice as the *Gnao1* mutant mice. *Gnao1* mutant male mice, generated from founders were crossed with female WT C57BL/6J mice from Jackson Labs (Bar Harbor, ME). Animals used for this study were between 5 and 10 weeks old. Only male heterozygous mutant animals were used due to the stronger behavioral phenotypes (Feng et al., 2019).

**Figure 1.**
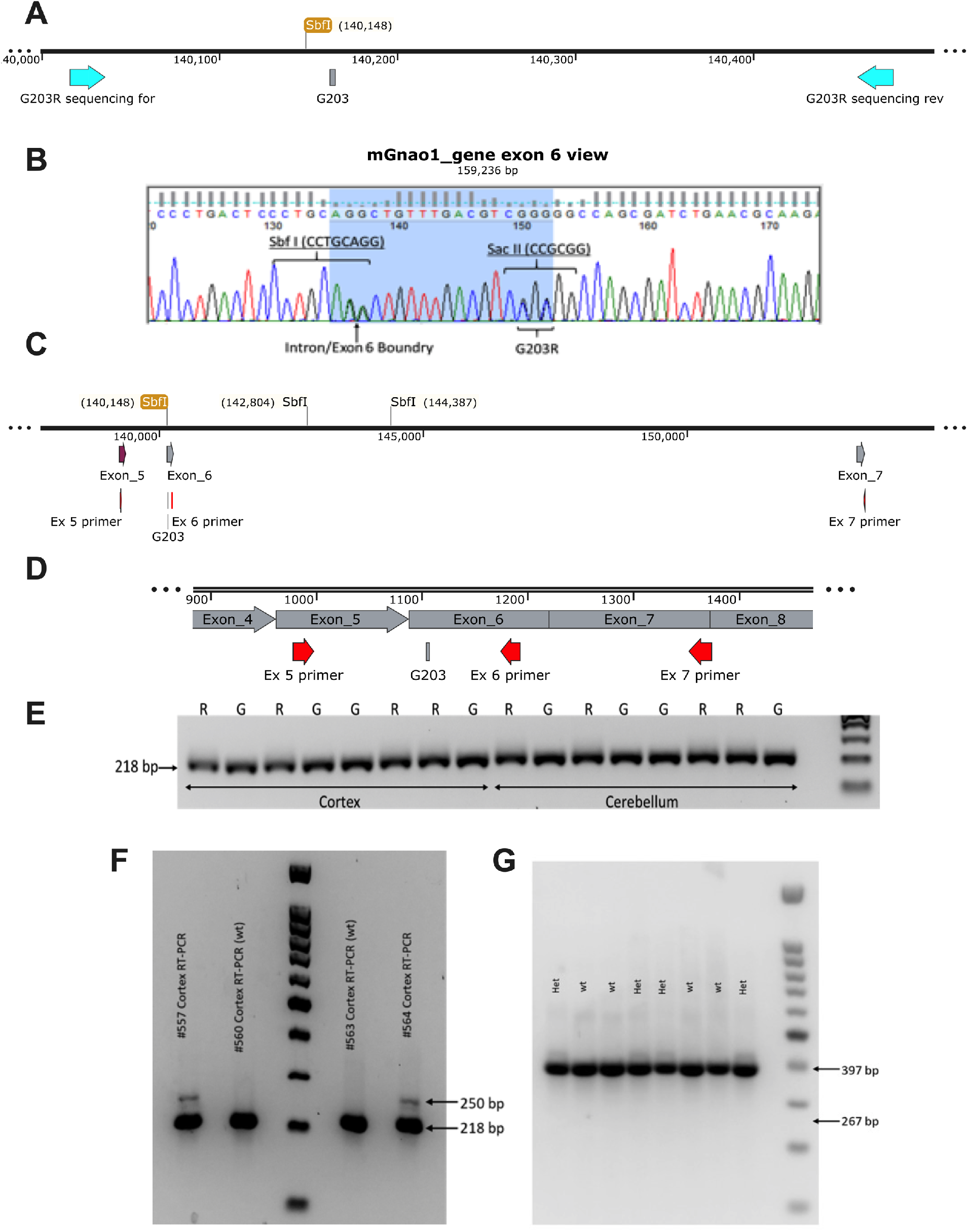
A second mutation in addition to G203R was identified in our previously described *Gnao1* mutant mouse model. (A) Genomic map around exon 6 showing the location of primers used for genotyping, the G203 codon, and an *SbfI* restriction site at the exon 6 splice acceptor site in the WT genomic sequence. (B) Sanger sequencing from a genomic PCR fragment around the G203R mutation in a heterozygous *Gnao1* mutant mouse (Feng et al., 2019). The expected G203R codon change (GGG > CGC) and an introduced *SacII* site are shown. An unexpected (heterozygous) sequence change (AGG > AAA) eliminates the *SbfI* site and also mutates the exon 6 splice acceptor sequence. A genomic (C) and mRNA (D) map of primers to assess splicing in RNA from brains of *Gnao1* mutant mice. (E) RT-PCR from cortex or cerebellum tissue of WT (gel lanes labelled G) and mutant (gel lanes labelled R) samples. A band of the expected size of 218 bp was seen for the RT-PCR product for the primers Ex 5 primer and Ex 6 primer. Sanger sequencing did not show any evidence of the G203R mutation in the spliced RNA product indicating that only the WT allele was appropriately spliced. (F) On a 2.5% agarose gel, a minor product of about 250 bp was observed only in Gnoa1 mutant mice. This is likely due to utilization of a weak splice acceptor site upstream of the usual Exon 6 splice acceptor but there was not sufficient product to show up with the G203R sequence in the Sanger traces. (G) RT-PCR using primers in Exons 5 and 7 showed only the expected band (397 bp) for the Exon 5+6+7 product with no evidence of a product with a skip over Exon 6 to splice Exon 5 and 7 directly.

### Tissue preparation and solutions

Mice were sacrificed by direct cervical dislocation and cerebellums were rapidly dissected. Parasagittal cerebellar slices (250 μm) were prepared using a Vibrotome™ 1000 (Leica, Wetzlar, Germany) according to previously described methods by Yuan and Atchison (Yuan & Atchison, 1999, 2003b, 2007, 2016). Before slicing, cerebellums were transferred into a chilled oxygenated sucrose-based slicing solution containing (in mM): 62.5, NaCl; 107, sucrose; 2.5, KCl; 5, MgCl_2_; 1.25, KH_2_PO_4_; 26, NaHCO_3_; 0.5, CaCl_2_ and 20, D-glucose (pH 7.35–7.4 when saturated with 95% O_2_ /5% CO_2_ at room temperature of 22–25°C). Slices were incubated in the oxygenated slicing solution for 15 min, then transferred into standard artificial cerebrospinal fluid (ACSF) solution at room temperature for at least 60 min before electrophysiological recordings. The standard ACSF contains (in mM): 125, NaCl; 2.5, KCl; 1, MgCl_2_; 1.25, KH_2_PO_4_; 26, NaHCO_3_; 2, CaCl_2_ and 20, D-glucose (pH 7.35–7.4 saturated with 95% O_2_/5% CO_2_ at room temperature).

### Electrophysiological recording

Whole-cell patch clamp recording methods were detailed in previous publications (Yuan & Atchison, 1999, 2003b, 2007, 2016). Slices were placed in a recording chamber and perfused with standard ACSF bubbled with 95% O_2_/5% CO_2_. Individual neurons were visualized with a Nomarski 40X water immersion lens with infrared differential interference contrast optics using a Nikon E600FN upright microscope (Nikon Inc., Chicago, IL). Recording electrodes were fire polished and had a resistance of 3–7 MΩ when filled with pipette solutions. For recording sIPSCs and mIPSCs, the pipette solution consisted of (in mM) 140, CsCl; 0.4, GTP; 2, Mg-ATP; 0.5, CaCl_2_; 5, Phosphocreatine Na_2_; 5, EGTA-CsOH; 10, HEPES (pH 7.3 adjusted with CsOH). For recording sEPSCs and mEPSCs, the pipette solution consisted of (in mM) 140, K-Gluconate; 4 NaCl; 0.4, GTP; 2, Mg-ATP; 0.5, CaCl_2_; 5, EGTA-CsOH; 10, HEPES (pH 7.3 adjusted with KOH). The holding potential was -70 mV for recording of both IPSCs and EPSCs. For recording inhibitory currents, 6-Cyano-7-nitroquinoxaline-2,3-dione (CNQX, 10 µM) and amino-5-phosphonopentanoic acid (APV, 100 µM) were added to the external solution to block glutamate receptor-mediated sEPSCs. For recordings of miniature IPSCs (mIPSCs), 0.5 μM tetrodotoxin (TTX) was added to the external solution in addition to CNQX and APV. For recording sEPSCs, bicuculline (10 µM) was added to the external solution to block GABA receptor-mediated sIPSCs. For recording of mEPSCs, the external solution was supplemented with 0.5 μM TTX in addition to bicuculline. Whole cell currents were filtered at 2–5 kHz with an 8-pole low-pass Bessel filter and digitized at 10–20 kHz for later off-line analysis using the pClamp 9.0 program (Molecular Devices, Inc., Sunnyvale, CA). All experiments were carried out at room temperature (22–25°C).

### Pharmacology

The following agents were used: CNQX disodium salts (Sigma-Aldrich, St. Louis, MO), APV acid solid (Sigma-Aldrich, St. Louis, MO), TTX (Tocris, Bristol, UK), pertussis toxin (PTX) (List Biological Laboratories, Campbell, CA), baclofen (Sigma-Aldrich, St. Louis, MO), NEM (Sigma-Aldrich, St. Louis, MO), UK14,304 (Sigma-Aldrich, St. Louis, MO), CGP36216 hydrochloride (Cayman Chemical, Ann Arbor, MI), cadmium chloride (Sigma-Aldrich, St. Louis, MO). All drugs were made up as 1000 x concentrated stock solutions in distilled water, aliquoted and stored at ∼20°C. Aliquots were thawed and dissolved in oxygenated ACSF immediately prior to use.

### SDS PAGE and Western Blots

Male mice (6-8 weeks old) were sacrificed, and their brains were dissected into different regions and flash-frozen in liquid nitrogen. For Western Blot analysis, tissues were thawed on ice and homogenized for 5 min with 0.5 mm zirconium beads in a Bullet Blender (Next Advance; Troy, NY) in RIPA buffer (20mM Tris-HCl, pH7.4, 150mM NaCl, 1mM EDTA, 1mM β-glycerophospate, 1% Triton X-100 and 0.1% SDS) with a protease inhibitor cocktail (Roche/1 tablet in 10 mL RIPA). Homogenates were centrifuged for 5 min at 4°C at 13,000 x G. Supernatants were collected and protein concentrations determined using the bicinchoninic acid method (BCA method; Pierce; Rockford, IL). Protein concentration was normalized for all tissues with RIPA buffer and 2x SDS sample buffer containing β-mercaptoethanol (Sigma-Aldrich, St. Louis, MO) was added. Thirty µg of protein was loaded onto a 12% Bis-Tris gel (homemade), and samples were separated for 1.5 hrs at 160V. Proteins were then transferred to an Immobilon-FL PVDF membrane (Millipore, Billerica, MA) on ice either for 2 h at 100 V, 400 mA or overnight at 30 V, 50 mA. Immediately after transfer, PDVF membranes were washed and blocked in Odyssey PBS blocking buffer (LI-COR Biosciences, Lincoln, NE) for 40 min at RT. The membranes were then incubated with anti-Gα_o_ (rabbit; 1:1,000; sc-387; Santa Cruz biotechnologies, Santa Cruz, CA) or anti-Gβ (recommended for detection of Gβ_1_, Gβ_2_, Gβ_3_ and Gβ_4_; mouse; 1:1000; sc-378; Santa Cruz biotechnologies, Santa Cruz, CA) and anti-actin (goat; 1:1,000; sc-1615; Santa Cruz) antibodies diluted in Odyssey blocking buffer with 0.1% Tween-20 overnight at 4°C. Following four 5-min washes in phosphate-buffered saline with 0.1% Tween-20 (PBS-T), the membrane was incubated for 1 hr at room temperature with secondary antibodies (1:10,000; IRDye® 800CW Donkey anti-rabbit; IRDye® 800CW Donkey anti-mouse; IRDye® 680RD Donkey anti-goat; LI-COR Biosciences) diluted in Odyssey blocking buffer with 0.1 % Tween-20. The membrane was subjected to four 5-min washes in PBS-T and a final rinse in PBS for 5 minutes. The membrane was kept in the dark and the infrared signals at 680 and 800nm were detected with an Odyssey Fc image system (LI-COR Biosciences). The Gα_o_ polyclonal antibody recognizes an epitope located between positions 90-140 of the Gα_o_ protein (Santa Cruz, personal communication).

#### Tissue preparation and solutions

For immunofluorescence experiments, cerebella were drop-fixed in 4% paraformaldehyde for 3 days at 4°C. Tissues were then cryoprotected in 30% sucrose for 2 days at 4°C before being frozen in Tissue-Tek O.C.T. Compound (Sakura Finetek USA, Inc., Torrance, CA). Parasagittal cerebellar sections were generated on a Cryostat (Fisher Scientific, Hampton, NH) at 30 µm. Tissue was stored free-floating in PBS and 0.1% sodium azide at 4°C

#### Immunofluorescence

For Kit-eGFP tissue, a dissection scope was used to identify medial slices of the cerebellar vermis corresponding to sections 20 & 21 of the sagittal Allen Mouse Brain Atlas. These sections were then mounted with ProLong Gold Antifade Mountant (Invitrogen, Waltham, MA) for imaging.

For immunohistochemistry, slices were allowed to warm to room temperature, rinsed in PBS, then permeabilized with PBS and 0.3% Triton X-100 for 30 minutes. Tissue was then blocked for 30 minutes in PBS plus 20% newborn calf serum (NBCS) and 0.3% Triton X-100. Slices were incubated in primary antibodies overnight at 4°C, rinsed with PBS, and incubated in secondary antibodies for 60 minutes at room temperature on a shaker. After being rinsed in PBS, slices were mounted as described. The following primary antibodies were used, diluted in PBS plus 2% NBCS and 0.3% Triton X-100: mouse anti-Glutamate decarboxylase (65 kD isoform, GAD65; 1:1500; ab26113; Abcam, Cambridge, United Kingdom), rabbit anti-Calbindin (1:3000; sysy-214 002; Synaptic Systems, Göttingen, Germany). Secondary antibodies were diluted similarly and include Alexa Fluor conjugated -488 and -546 (1:500; Invitrogen, Waltham, MA).

#### Kit-eGFP labeling of molecular layer inhibitory neurons

In postnatal mice, Kit is expressed at the transcript and protein level in GABAergic molecular layer interneurons (MLIs) of the cerebellar cortex. To visualize MLIs, we utilized a previously established BAC transgenic mouse line harboring an eGFP-polyA sequence regulated by the Kit locus (Heintz, 2004). In both male and female mice, and across all generations of backcrossing to a C57BL/6J background, we found that the pattern of GFP expression matched the literature for Kit expression in the cerebellar cortex: expression in the MLIs, and sparse expression in Golgi cells (Amat et al., 2017; Hirota et al., 1992; Keshet et al., 1991; Manova et al., 1992; Motro, van der Kooy, Rossant, Reith, & Bernstein, 1991). The Kit eGFP strain used in this study was generated from (Tg(Kit-EGFP)IF44Gsat/Mmucd; Gensat) (Heintz, 2004) that had been backcrossed for greater than 5 generations to C57BL6/J (Stock No: 000664, The Jackson Laboratory). These mice were further crossed with heterozygous *Gnao1* mutant mice to generate Kit-eGFP expressing littermates with WT or LOF *Gnao1* alleles.

#### Image Analysis

Epifluorescent images were collected with a Nikon Eclipse Ti2-E Inverted Microscope (Nikon Inc., Chicago, IL) at 20X magnification. Images shown are Z stacks of the entire focus plane at a step size of 0.9 μm. True confocal images were captured on an Olympus FV1000 confocal laser scanning microscope (Olympus America Inc., Center Valley, PA) with a 63X oil-immersion objective and a step size of 0.43 μm. Z stacks of 14 slices were collected to capture the pinceau and perisoma of multiple PCs. All images were edited in the Fiji distribution of ImageJ2 (Schindelin et al., 2012). GAD65 and 40X Kit-eGFP images were deconvolved using the DeconvolutionLab plugin and the Richardson-Lucy algorithm, 10 iterations. Point-spread functions were generated using the PSF Generator plugin with the Born & Wolf 3D Optical Model (Kirshner, Aguet, Sage, & Unser, 2013).

### Statistical Analysis

Electrophysiological data analysis was performed as described previously (Yuan & Atchison, 1999, 2003b, 2007, 2016). The individual performing the analysis was blinded to the genotype until all results were recorded. In brief, spontaneous synaptic currents were first screened automatically using MiniAnalysis software (Synaptosoft Inc., Decatur, GA) with a set of pre-specified parameters (i.e. period to search a local maximum; time before a peak for baseline; period to search a decay time; fraction of peak to find a decay time; period to average a baseline; area threshold; number of points to average for peak; direction of peak). They were accepted or rejected manually with an event detection amplitude threshold at 5 pA for sIPSCs/mIPSCs and 3 pA for sEPSCs/mEPSCs as well as being assessed for the typical kinetic properties (fast rising phase and slow decay phase) of the spontaneous events. These thresholds were chosen based on the baseline noise level of each individual recording (baseline noise amplitude ranged from to 0.8 pA) so the thresholds were from 6-25x the noise level. Unless otherwise specified, synaptic events per cell collected over a 2-min period were averaged to calculate the frequency and amplitude of spontaneous synaptic currents. Amplitudes of currents were measured after subtraction of the baseline noise. MiniAnalysis-derived results were plotted in GraphPad Prism (GraphPad; LaJolla, CA). Results from more than one neuron from a single animal were averaged prior to statistical analysis. Some graphs, when indicated, show points for each individual neuron while bar graphs and error bars are calculated from the per animal data. Data are presented as mean value ± SEM, where n=number of animals. Statistical significance was determined using unpaired Student’s t-test unless stated otherwise. A p value < 0.05 was deemed significant.

Quantification of infrared (IR) Western blot signals was performed using Image Studio Lite (LI-COR Biosciences). Individual bands were normalized to the corresponding actin signals, and WT Gα_o_ was set as control for each blot. All data were analyzed using GraphPad Prism 8.0 (GraphPad; LaJolla, CA).

For cell counts, images were thresholded to mask cells within the focus plane. The watershed plugin in ImageJ2 was used to separate overlapping cells. The *Analyze Particles* function was used to count cells and measure their sizes. To avoid artifact, the average intensity of particles measured during cell counts were compared between WT and *Gnao1* group, no significant difference was observed. Pinceau were traced manually from Z-projections. Perisomatic contacts with basket cell axons were assessed by measuring the thickness of Kit-eGFP fluorescence at each PC’s widest point.

GAD65 staining in the molecular layer was also quantified using the *Analyze Particles* function. Measurements were taken in a rectangular ROI above the PC layer on a projection of 3 Z-steps. To exclude off-target vasculature staining, particles larger than 3 µm^2^ were not measured. Pinceau were measured on a projection of 3 Z-steps at each PC’s widest point. The magic wand tool was used to define each ROI. GAD65 puncta on the periphery of each PC were manually counted to assess perisomatic synapses. For validation, the cell counts were repeated at a different magnification using a different method (auto-thresholding vs. manual).

## Results

### Identification of second site mutation

C57BL/6J mice containing a G203R mutant allele were generated as described (Feng et al, 2019) where the mutant allele introduces a new Sac II restriction site which was used for genotyping. Subsequently, a tail snip from a heterozygous G203R mutant mouse was obtained for genomic PCR and Sanger sequencing to define the local DNA sequence around the mutation. In addition to the G203R mutation, there were two additional heterozygous base alterations just upstream of the designed mutation (Figure 1). This resulted in the sequence AAA replacing the AGG at the exon 6 splice acceptor site. The second mutation (AGG>GAA) destroys an Sbf1 restriction site that is present in the wild-type sequence. To determine whether this second mutation was a random mutation present in a single mutant mouse, we used genomic PCR and restriction digestion with SacII (G203R) and SbfI (splice site) and screened 52 mice from 10 litters. All 30 mice carrying the G203R mutation also had loss of the Sbf1 site and none of the 22 mice with the WT codon 203 sequence had the loss of Sbf1 digestion.

Review of older data from the founder mouse revealed that the second mutation was present at the initial development of the mouse line. The way the sequence data were collected (sequencing from a cloned PCR product) caused us to miss the fact that the sequence did not fully match the WT sequence. Thus all G203R mutant mice in this present study and in our previous publication (Feng et al., 2019) appear to have the second site mutation at the splice site.

### Loss of splice site activity

Introns nearly universally terminate in an AG base pair at the splice acceptor site. Since the second mutation converted an AGG to AAA it is highly likely to disrupt normal splicing of Exon 5 onto Exon 6. To determine if this was the case, we undertook RT-PCR with primers in exons 5 and 6 and in exons 5 and 7 to understand how splicing is perturbed by the second site mutation. RT-PCR of brain tissue from heterozygous G203R mutant mice using primers in exons 5 and 6 yielded a 218 bp band of the expected size for normal splicing. Sequence analysis, however, showed only the WT sequence at codon 203. Therefore, there was no normal splicing in the G203R mutants. Those samples did reveal a faint band of higher molecular weight, which we isolated and sequenced, that showed a rare splicing event 45 bp upstream of the usual splice junction that occurred only in the G203R mutants (Figure 1). The sequence of this alternatively spliced cDNA showed a stop codon before codon 203.

RT-PCR using primers in exons 5 and 7 also revealed bands of the appropriate size 397 b.p. There was no evidence of skipping of exon 6 to produce a direct splice from exon 5 to exon 7 (expected size 267 b.p.). The low frequency insertion of 45 bp seen with the exon 5 and 6 primers was also evident in the samples from mutant mice for the exon 5-7 RT-PCR (Figure 1E).

### Gnao1 mutant mice exhibit decreased Gα_o_ protein but no change in Gβ levels

We tested Gα_o_ protein expression in whole brain and selected brain regions of WT and *Gnao1* mice. Results showed a significant reduction in Gα_o_ protein expression (∼50% of WT) in whole brain (Figure 2A, 2C; p<0.0001). The decrease in Gα_o_ protein expression was also significant in cerebellum, cortex, hippocampus, and striatum of the mutant mice (Figure 2B, 2D; p<0.05). No significant change was observed in the brainstem or the olfactory bulb.

**Figure 2.**
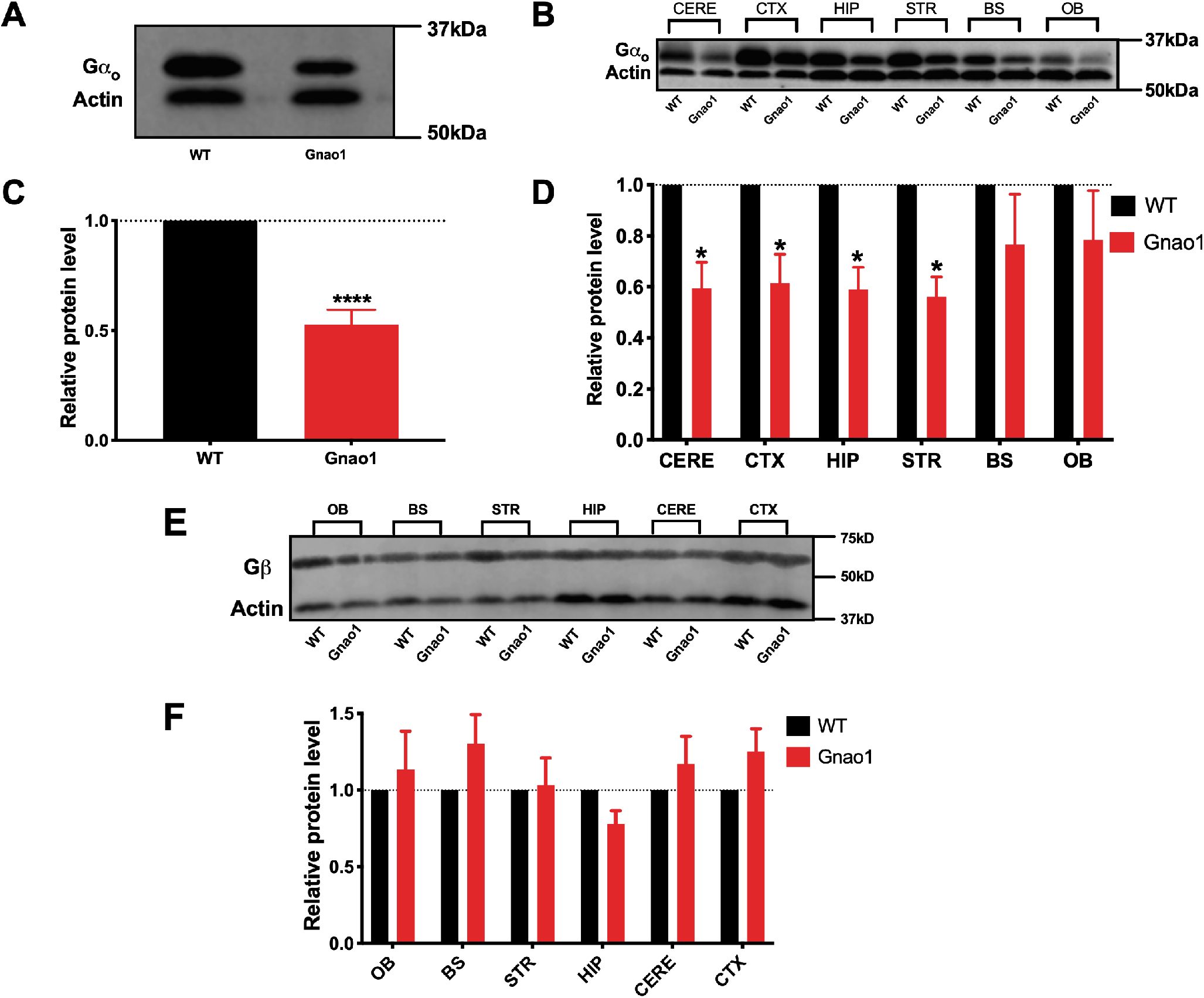
*Gnao1* mice showed a significant decrease in Gα_o_ protein expression but normal Gβ expression level. (A, C) Whole brain Gα_o_ expression levels were about 50% of WT in mutant *Gnao1* mouse brain lysates. (B, D) This reduction in expression was most significant in cerebellum (CERE), cortex (CTX), hippocampus (HIP) and striatum (STR), while expression was not significantly reduced in brain stem (BS) and olfactory bulb (OB). All expression levels were normalized to that of WT. Unpaired Student’s t-test; WT (n=8), *Gnao1* (n=8); ****p<0.0001, *p<0.05. (E) A representative gel shows the Gβ protein expression patterns in each individual brain region. (F) Quantification of Gβ protein expression levels is unchanged in *Gnao1* mice brain lysates comparing to those of their WT siblings. Unpaired Student’s t-test; WT (n=4), *Gnao1* (n=4).

Interestingly, expression levels of Gβ did not change in *Gnao1* mutant mice (Figure 2E & 2F). Given that Gα_o_ is the most abundant Gα subunit in brain, a decreased Gα_o_ protein expression level accompanied by normal Gβ protein levels suggests that there would be increased free Gβγ subunits, which could lead to enhanced receptor-independent Gβγ signaling.

### Presynaptic GABA release is suppressed in cerebellar PCs of Gnao1 mutant mice

sIPSCs, which are considered to be due to AP-dependent GABA release, were isolated by adding 10 μM CNQX and 100 μM APV in standard ACSF to block glutamate signaling. sIPSC frequency was decreased by 40% in *Gnao1* mice compared to their WT siblings (WT: 21.0 ± 1.7 Hz; *Gnao1*: 12.7 ± 2.5 Hz; Figure 3A & 3C, p<0.01), but there was no difference in sIPSC amplitude (WT: 41.8 ± 8.5 pA; *Gnao1*: 36.7 ± 7.4 pA; Figure 3B & 3D, n.s.), suggesting a presynaptic mechanism.

**Figure 3.**
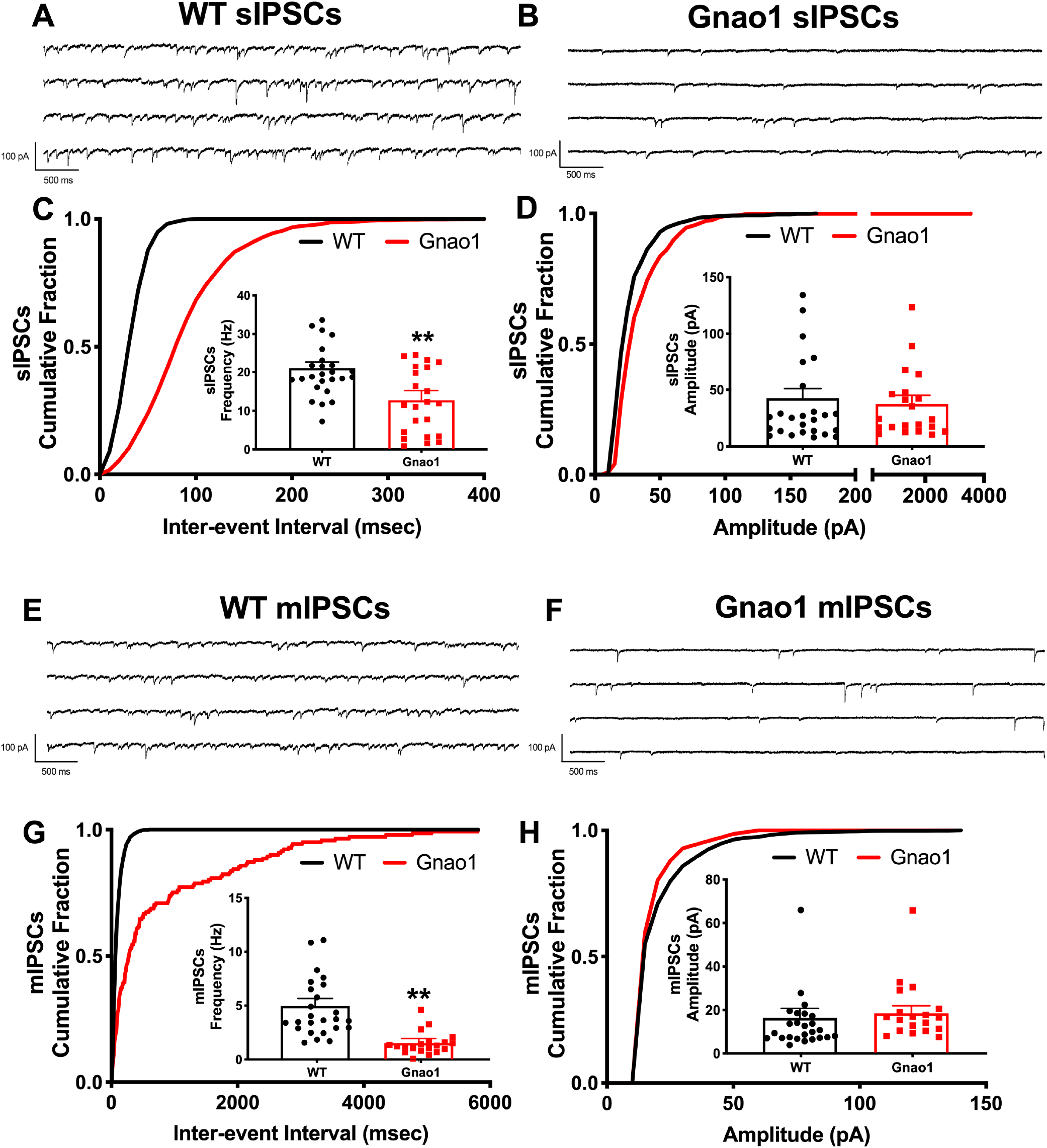
Cerebellar Purkinje cells in brain slices from 4-6 week-old *Gnao1* mutant mice display reduced GABAergic spontaneous synaptic currents (sIPSCs) and reduced miniature synaptic currents (mIPSCs). (A, B) Representative recording of spontaneous inhibitory postsynaptic currents in a cerebellar Purkinje cell from a 4 week-old mouse in the presence of 10 μM of CNQX and 100 μM of AP-V at a holding potential of -70 mV. (C) *Gnao1* mice showed a decrease in the frequency of sIPSCs. (D) No significant difference is observed in the amplitude of sIPSCs between WT and *Gnao1* mice. Unpaired Student’s t-test; **p=0.0086 WT (n=13 mice), *Gnao1* (n=9 mice). (E, F) Representative recording of spontaneous miniature inhibitory postsynaptic currents in a cerebellar Purkinje cell from a 4 week-old mouse in the presence of 10 μM of CNQX, 100 μM of AP-V and 0.5 μM TTX at a holding potential of -70mV. (G) *Gnao1* mice showed a decrease in the frequency of mIPSCs. (H) No significant difference is observed in the amplitude of mIPSCs between WT and *Gnao1* mice. Unpaired Student’s t-test; **p=0.0011; WT (n=13 mice), *Gnao1* (n=9 mice). Recordings from each cell are shown as a data point but the bar graph, error bars, and statistical analysis was averaged data per animal. Data were recorded from 25 cells of 13 WT mice for WT and 21 cells from 9 *Gnao1* mice.

mIPSCs, which are due to AP-independent GABA release, were isolated by the further addition of 0.5 μM TTX. As reported previously in cerebellar PCs, TTX significantly reduced mean sIPSC frequency and amplitude (Bardo, Robertson, & Stephens, 2002; Boxall, 2000; Harvey & Stephens, 2004; Yuan & Atchison, 2003a). Slices from *Gnao1* mice exhibited a marked 75% reduction in mIPSC frequency compared to those of WT mice (Figure 3E & 3G, p<0.01), but not in mIPSC amplitude (Figure 3F & 3H, n.s.). The percentage change in mIPSC frequency was greater for mIPSCs than that for sIPSCs (75% vs 40% decrease).

### Presynaptic glutamate release is not affected by the LOF Gα_o_ protein

To investigate whether the LOF mutant affects the excitatory inputs to PCs, we recorded sEPSCs and mEPSCs from PCs. The AP-dependent sEPSCs were isolated with 10 μM bicuculline in the bath and AP-independent mEPSCs were recorded with the further addition of 0.5 μM TTX. Although the mutant Gα_o_ protein significantly decreased sIPSC and mIPSC frequency, EPSCs are seemingly affected in neither frequency nor amplitude for both sEPSCs and mEPSCs (Figure 4; sEPSC frequency: WT: 1.50 ± 0.20 Hz vs. *Gnao1*: 1.36 ± 0.43 Hz, n.s.; sEPSC amplitude: WT: 7.52 ± 0.92 pA vs. *Gnao1*: 6.42 ± 0.50 pA, n.s.; mEPSC frequency: WT: 1.05 ± 0.16 Hz vs. *Gnao1*: 0.99 ± 0.46 Hz, n.s.; mEPSC amplitude: WT: 6.39 ± 0.91 pA; *Gnao1*: 6.24 ± 0.60 pA, n.s.).

**Figure 4.**
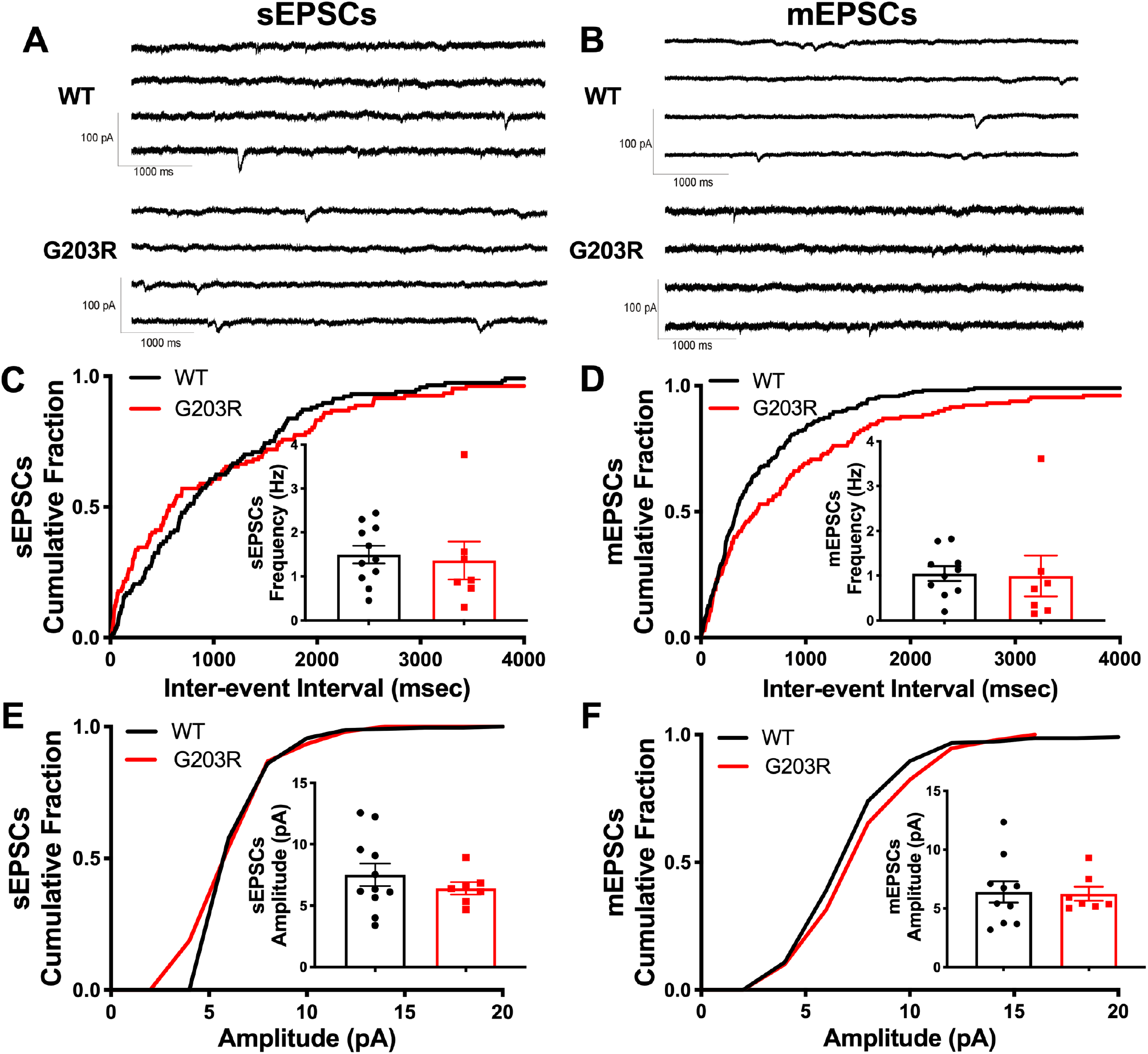
*Gnao1* mutant slices show no difference in either sEPSCs or mEPSCs. (A) sEPCSs were recorded from Purkinje cells at a holding potential of -70mV in the presence of 10 μM bicuculline. (B) 0.5 μM TTX was then added to the bath in order to record mEPSCs. (C, E) No significant difference in either frequency or amplitude was observed between WT and *Gnao1* mice. (D, F) No significant difference between WT and *Gnao1* mice was observed in mEPSCs either. Unpaired Student’s t-test; WT (n=5), *Gnao1* (n=5). Data were recorded from 11 cells of 5 WT mice for WT and 6 cells from 5 *Gnao1* mice.

### Pertussis-toxin reverses the enhanced inhibition of mIPSC frequency in Gnao1 mice

Considering the previously proposed GOF mechanism of the G203R mutant Gα_o_, we examined the effects of PTX incubation on the AP-dependent and AP-independent IPSCs in cerebellar PCs. PTX catalyzes the ADP-ribosylation of α subunits of the heterotrimeric G_i/o_ family, thereby preventing the G proteins from interacting with GPCRs (Mangmool & Kurose, 2011). Slices in this experiment were subject to incubation in ACSF containing 1 μg/mL PTX for more than 6 hours before recording. Surprisingly, PTX significantly increased the mIPSC frequency in slices from *Gnao1* mice but had no effect on WT mice (WT: from 4.10 ± 0.70 Hz to 3.40 ± 0.68 Hz, n.s.; *Gnao1*: from 1.31 ± 0.16 Hz to 2.29 ± 0.30 Hz, p<0.01; Figure 5B & 5D). The mean amplitude of mIPSCs was not changed in either WT or *Gnao1* mice after PTX incubation (WT: 12.1 ± 1.6 pA to 9.7 ± 1.4 pA, n.s., *Gnao1*: 19.6 ± 1.6 pA to 15.8 ± 2.9 pA, n.s.; Figure 5C & 5E). In contrast to mIPSCs, sIPSC frequency was not affected by PTX in either WT or *Gnao1* mice (Frequency: WT: 13.8 ± 1.7 Hz to 10.9 ± 3.6 Hz, n.s.; *Gnao1*: 9.1 ± 1.4 Hz to 12.5 ± 2.3 Hz, n.s.; Amplitude: WT: 18.9 ± 3.4 pA to 19.2 ± 8.9 pA, n.s.; *Gnao1*: 29.2 ± 4.1 pA to 24.4 ± 4.5 pA, n.s.; Figure 5-1). Given the reduced expression of Gα_o_ in the *Gnao1* mutant mice, the effect of PTX to reverse the inhibition of mIPSCs is likely indirect (see Discussion).

**Figure 5.**
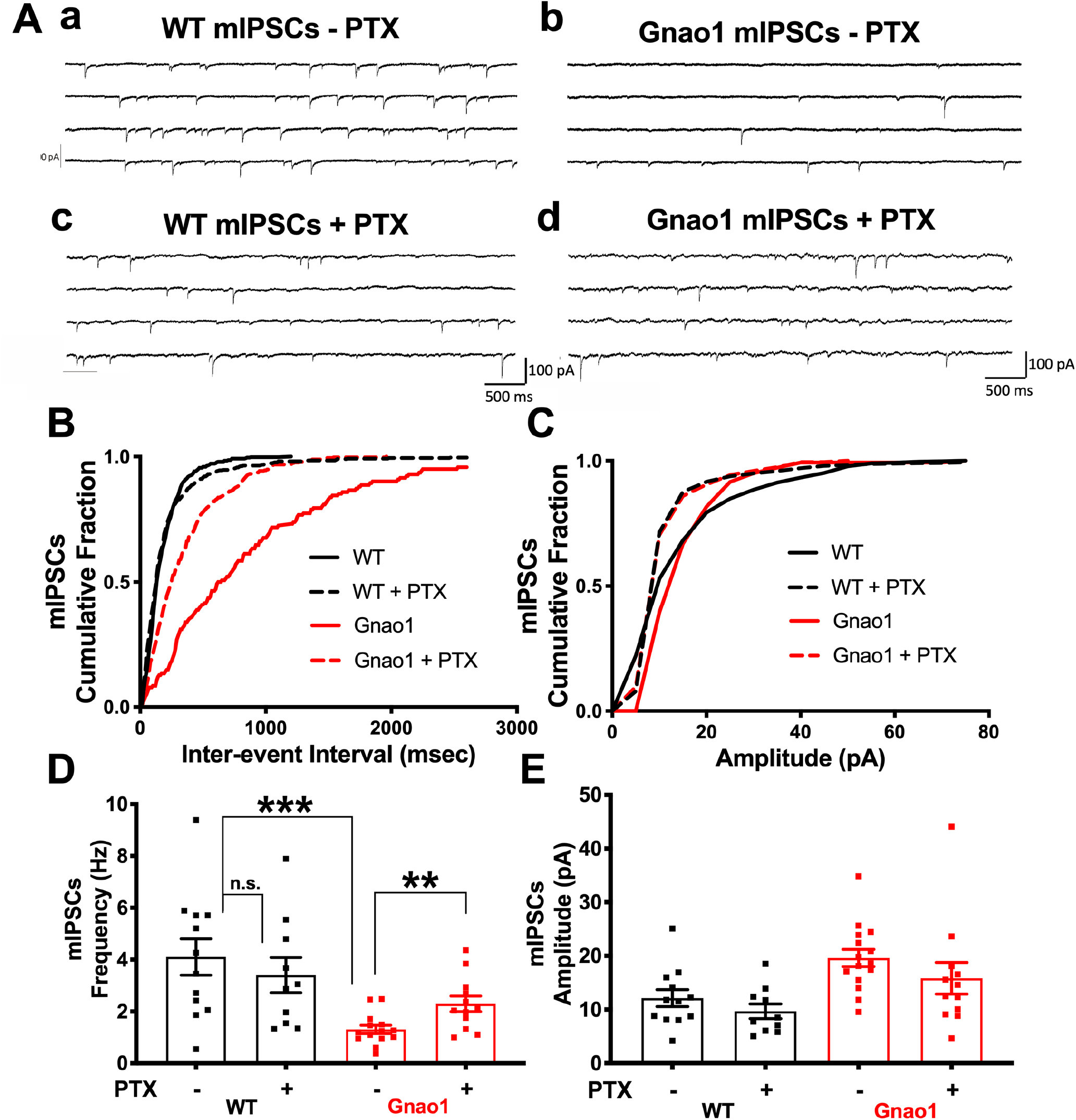
A selective inhibitor of G_i/o_, PTX, increased the frequency of mIPSCs in *Gnao1* but not WT mice. Slices were incubated in 1 μM/ml of PTX for >6 hrs pre-recording. Representative traces showed the example recordings of (A) sIPSCs in WT and *Gnao1* mice before and after PTX incubation. (B, D) PTX incubation significantly relived the G_o_ mediated inhibition of mIPSC frequency in *Gnao1* mice, but not WT mice. Unpaired Student’s t-test; WT (n=5 mice), *Gnao1* (n=6 mice); **p=0.006. (C, E) PTX did not change the mIPSC amplitude of either *Gnao1* or WT mice. Data were recorded from 11 cells of 5 WT mice and 12 cells from 6 *Gnao1* mice.

### Effects of GABA_B_ receptors on AP-independent GABA release onto PCs

To explore the effects of GABA_B_ receptors on reduced GABA release and to determine whether GABA_B_ responses are altered in the *Gnao1* mutant mice, mIPSCs were recorded in the presence of baclofen, a selective GABA_B_ receptor agonist. After recording baseline mIPSCs, baclofen (10 μM) was applied to the bath. Baclofen caused a clear reduction in mean mIPSC frequency in slices from both WT (from 5.47 ± 0.80 Hz to 1.19 ± 0.25 Hz, 78% inhibition, p<0.001) and *Gnao1* mice (from 1.24 ± 0.20 Hz to 0.65 ± 0.11 Hz, 48% inhibition, p<0.05) after 4 to 8 min (Figure 6C & 6E). Mean mIPSC amplitude was unchanged by baclofen (Figure 6D & 6F; WT: 12.8 ± 2.3 pA to 13.6 ± 4.3 pA, n.s.; *Gnao1*: 17.8 ± 1.9 pA to 17.3 ± 1.6 pA, n.s.), consistent with a presynaptic mechanism of action. Interestingly, the application of baclofen did not affect sIPSCs, while α_2_ adrenergic receptors only regulate sIPSCs but not mIPSCs (Harvey & Stephens, 2004; data not shown).

**Figure 6.**
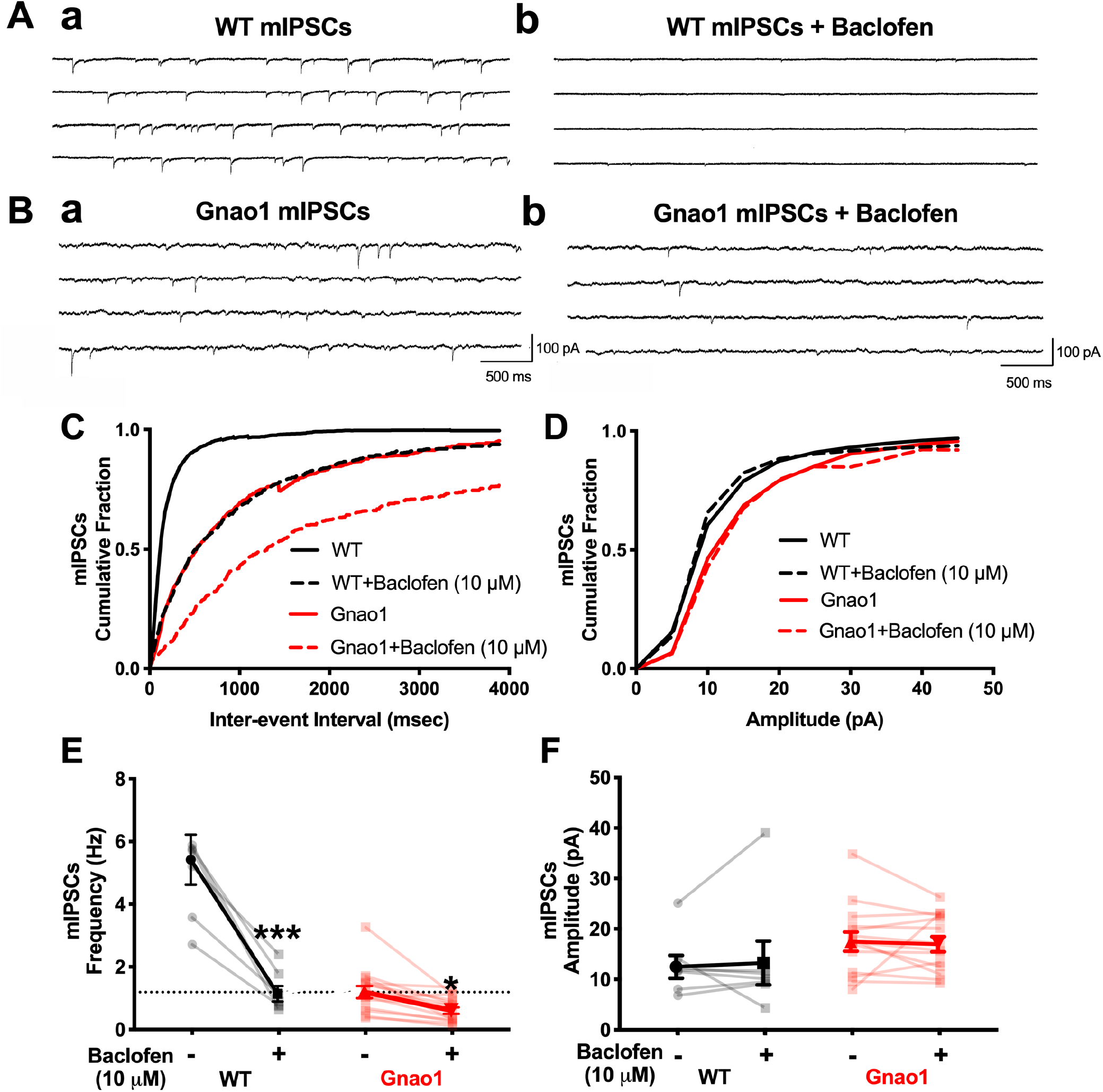
Activating GABA_B_ receptor with baclofen reduces mIPSC frequency but not amplitude. Representative traces showing the reduced mIPSC responses before and after adding baclofen (10 μM) in WT (A) and *Gnao1* (B) mice w/o baclofen. (C, E, G) Baclofen significantly decreased the frequency of mIPSC and the difference between WT and *Gnao1* remains though baclofen was present. Unpaired Student’s t-test; WT (n=6), *Gnao1* (n=6); ****p<0.001, ***p=0.003, *p=0.029 (WT vs. *Gnao1*), *p=0.014 (*Gnao1* w/o baclofen). (D, F, H) Amplitude remained unchanged regardless of the existence of baclofen.

G_o_ or related G_i_ proteins contribute to the baclofen-induced inhibition of mIPSC frequency as previously reported (Harvey & Stephens, 2004). PTX (1 μg/mL) eliminated baclofen-induced inhibition of mIPSC frequency (Figure 6-1 C, E). PTX did not affect mIPSC amplitude (Figure 6-2 D, F). Cadmium Cd^2+^, which blocks calcium channels, did not affect the frequency or amplitudes of mIPSCs in either WT or *Gnao1* mice (Figure 6-2) in contrast to the effect of PTX (Figure 5). The baclofen-induced decrease of mIPSC frequency was also unaffected by Cd^2+^ (Figure 6-2) which implies direct synaptic inhibition rather than voltage-gated calcium channel effects as the mechanism.

### No significant difference in the baclofen concentration-response curve between WT and Gnao1 mutant

We tested the baclofen concentration dependence in reducing PC mIPSCs. Baclofen (1, 3, and 10 μM) was added cumulatively into the ACSF while recording mIPSCs. Higher concentrations of baclofen inhibited the mIPSC frequency to a greater extent. However, there was no significant difference in the baclofen concentration-response between WT and *Gnao1* when the results were normalized to the baseline mIPSC level without baclofen (Figure 7). This observation suggests that the difference in mIPSC frequency caused by the *Gnao1* mutant is independent of the activation of GABA_B_R, which would be expected for a signal caused by free Gβγ due to reduced Gα_o_ levels.

**Figure 7.**
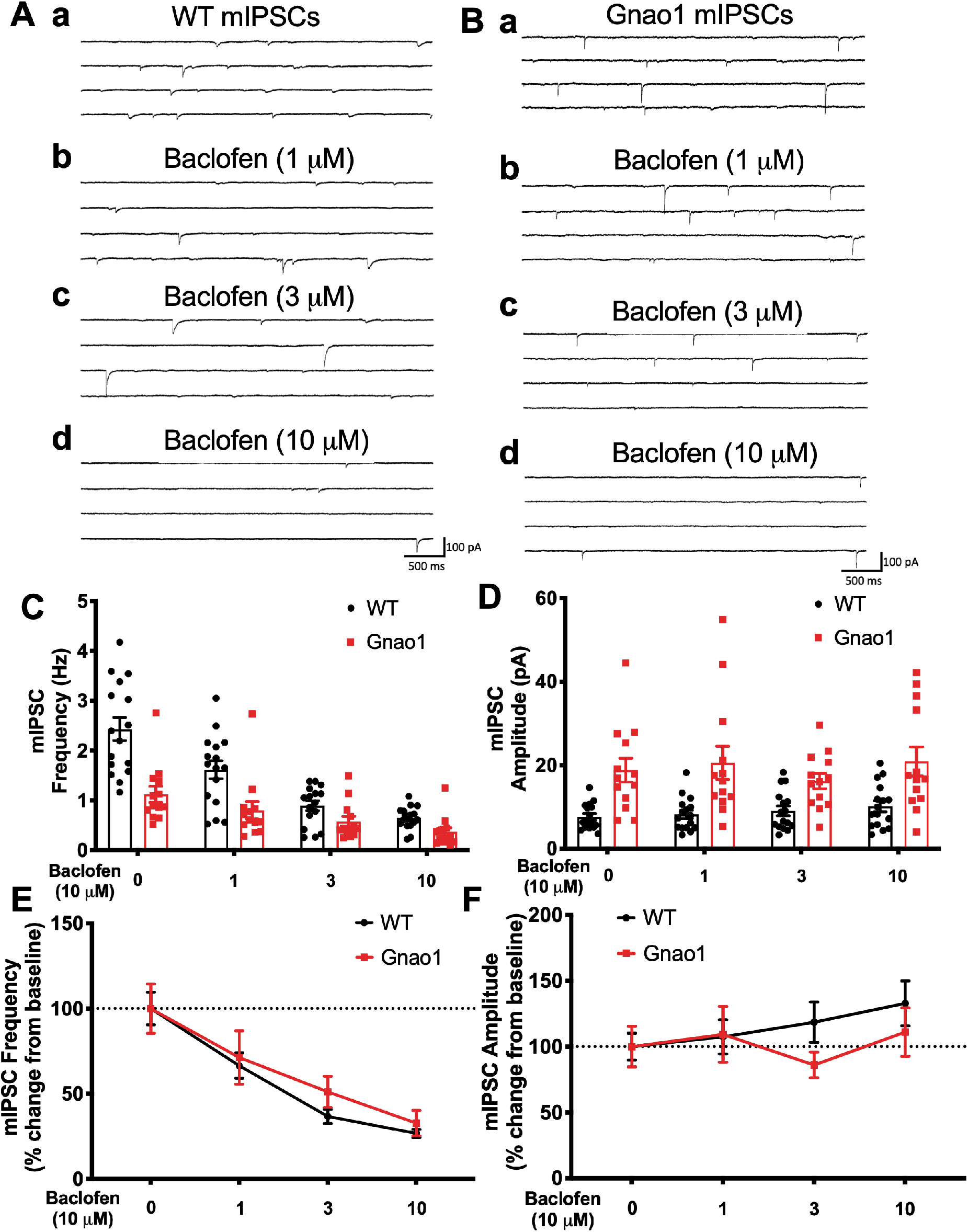
Baclofen dose-response curves of WT and *Gnao1* are similar to each other. Representative traces showing the mIPSC recording with no baclofen (a), 1 μM baclofen (b), 3 μM baclofen (c), 10 μM baclofen (d) in WT (A) and *Gnao1* (B) mice. Bar graph of mIPSC frequency (C) and amplitude (D) w/o baclofen with WT and *Gnao1* mice showed similar trend of baclofen-induced inhibition of mIPSC frequency but not amplitude. Each data point was normalized to the baseline level (no baclofen), and there was no significant dose-response curve shift in either frequency (E) or amplitude (F). WT (n=10), *Gnao1* (n=8).

### Pre-treatment with GABA_B_R antagonist CGP36216 (100 μM) prevented baclofen-induced inhibition of mIPSC frequency but did not reverse Gnao1 mutant suppression of mIPSC frequency

Pre-treating PCs with the GABA_B_R antagonist CGP36216 (100 μM) did not enhance baseline mIPSC frequencies in either WT or *Gnao1* cerebellar slices (Figure 8C). It did, however, eliminate the baclofen-induced inhibition of mIPSC frequency (Figure 8C). mIPSC amplitude was not affected (Figure 8D). The lack of effect of a GABA_B_R antagonist on baseline mIPSCs was observed previously with another GABA_B_R antagonist (Harvey & Stephens, 2004) and suggests that there is no tonic regulation of GABA release by GABA_B_ receptors in WT mice. Surprisingly, the same is true for the *Gnao1* mutant mice, which have a suppressed mIPSC frequency, further supporting the independence of the effect of *Gnao1* mutant from GABA_B_ receptors. These data are consistent with a presynaptic role of G_o/i_ subunits in baclofen-induced inhibition of AP-independent GABA release onto PCs by a mechanism that is also independent of membrane calcium channels. The direct Gβγ-mediated regulation of synaptic vesicle fusion (Blackmer et al., 2001; Zurawski et al., 2017) seems to be the most likely explanation.

**Figure 8.**
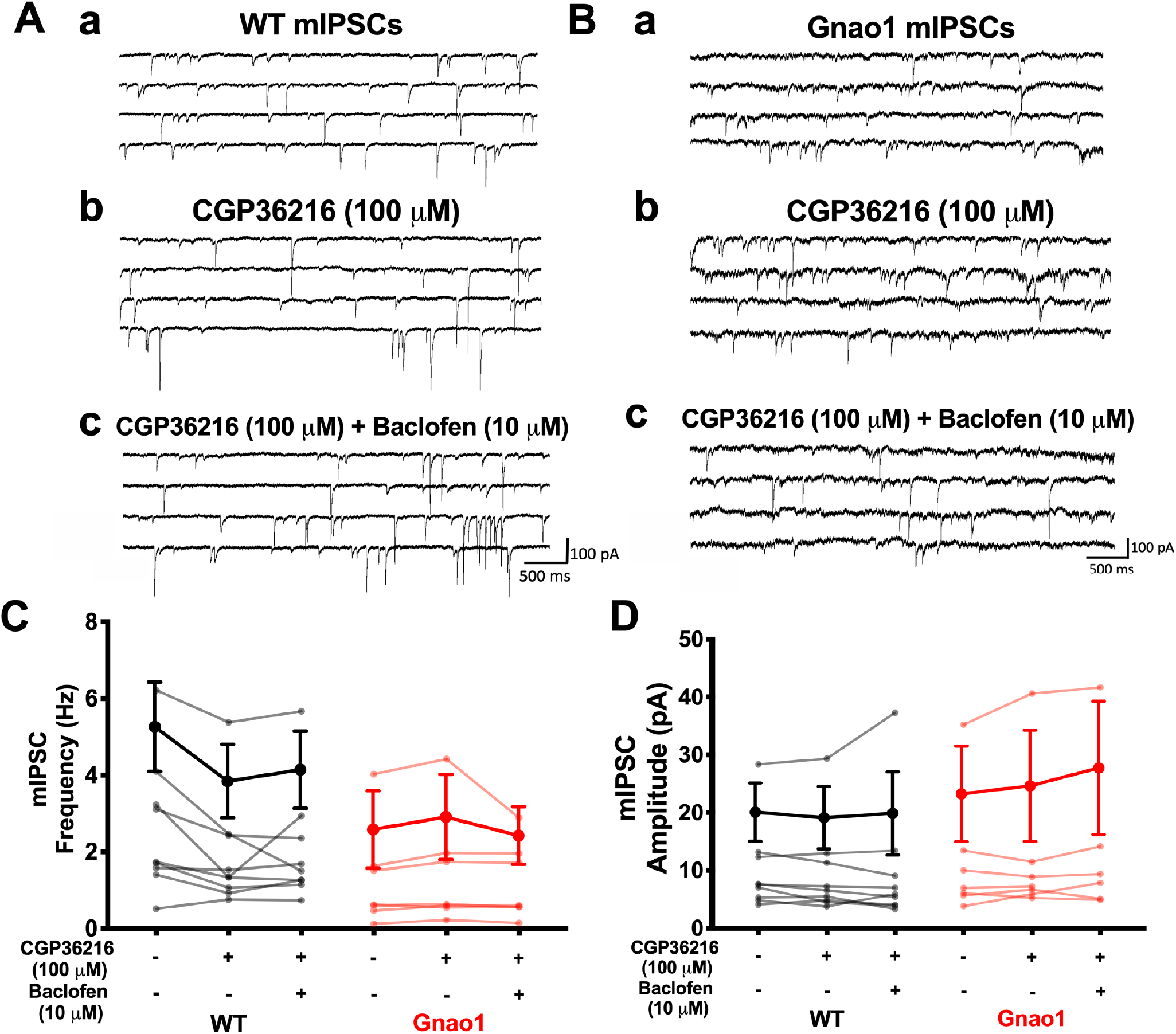
Pre-treatment of CGP36216 (100 μM) abolished baclofen (10 μM) - induced inhibition of mIPSC frequency. Representative traces showing no significant difference between baseline mIPSC levels (a) of WT and *Gnao1* with CGP36216 (b; 100 μM) and baclofen (c; 10 μM). CGP36216 (100 μM) eliminated the baclofen-induced decrease of mIPSC frequency in both WT and *Gnao1* (C). Amplitudes remained unchanged despite the drug treatment (D). WT (n=7), *Gnao1* (n=7).

### The number of cerebellar molecular layer interneurons is modestly reduced in Gnao1 mutant mice

Analysis of a c-kit eGFP reporter mouse revealed that interneuron number was significantly lower in cerebellar molecular layer (Figure 9). Cerebellar molecular layer mainly contains basket and stellate interneurons, which are important controllers of cerebellar cortical output by controlling the inhibitory synaptic input to PCs (Brown et al., 2019). Although there were fewer interneurons in *Gnao1* mutant’s cerebellar molecular layer, synapse count or axons contacts around PCs from basket and stellate cells remained unchanged (Figure 9-1 and 9-2).

**Figure 9.**
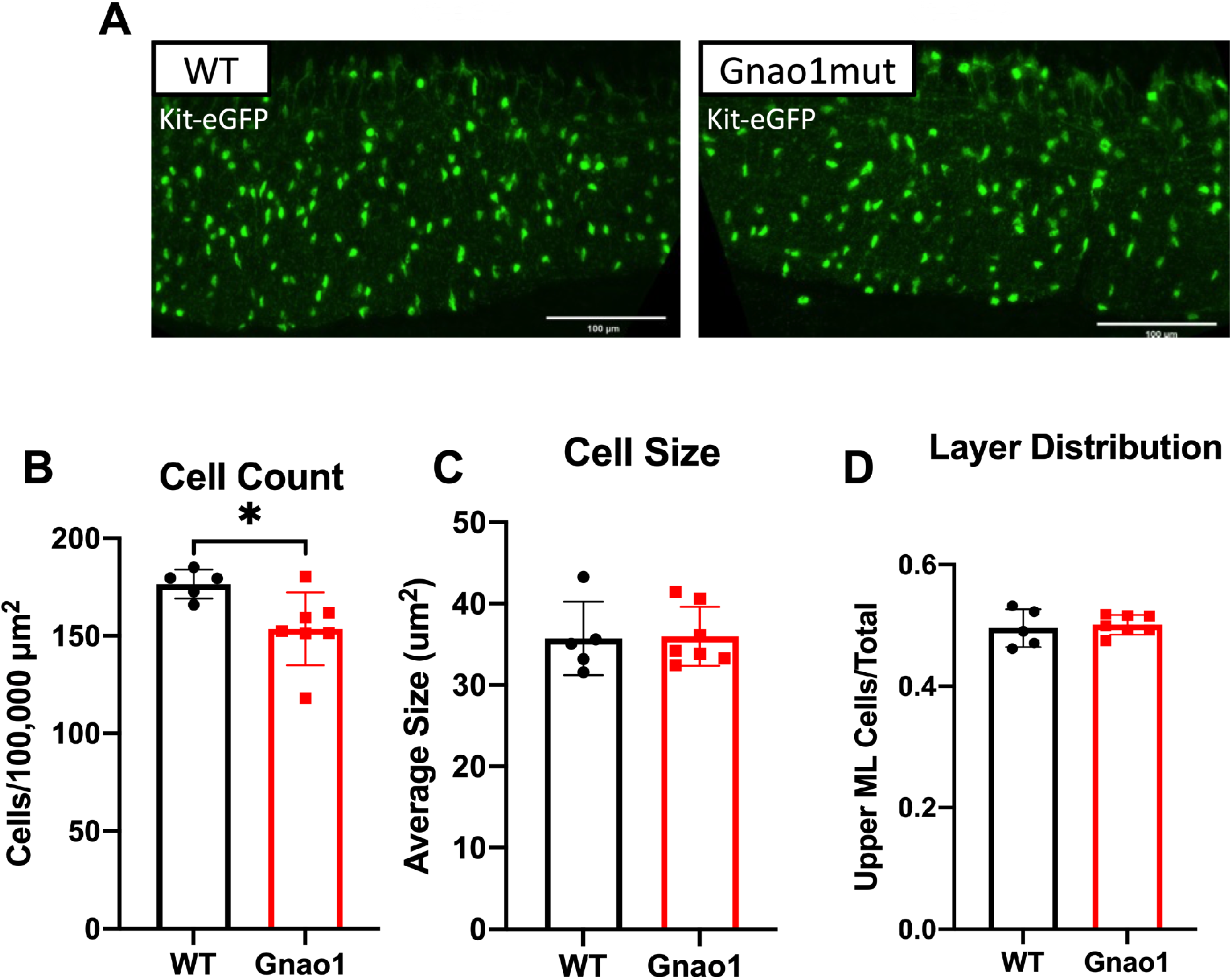
*Gnao1* mutants have a modest reduction in molecular layer interneuron count. (A) Representative immunofluorescent images of WT Kit-eGFP and *Gnao1* mutant x Kit-eGFP cerebellar cortex, lobe III. (B,C) *Gnao1* mutant mice have significantly fewer but not smaller molecular layer interneurons than WT. Unpaired Student’s t-test; WT (n=5), Gnao1 (n=7); *p=0.027 (count). (D) No significant difference in the distribution of cells between the upper and lower molecular layers was observed between WT and *Gnao1* mutant mice, suggesting a normal distribution of stellate and basket cells in mutants.

## Discussion

Mutations in Gα_o_ result in *GNAO1* encephalopathies which include severe epilepsy (Nakamura et al., 2013) or movement disorders (Ananth et al., 2016; Feng, Khalil, Neubig, & Sidiropoulos, 2018; Saitsu et al., 2016) with profound impacts on affected children. While cell-based studies (Feng et al., 2018; Feng et al., 2017) and animal behavioral models (Feng et al., 2019; Larrivee et al., 2019) have provided some mechanistic information about the human *GNAO1* mutations, there have been no neurophysiological studies of effects on synaptic transmission. Here, we show that a LOF mutant mouse with haploinsufficient Gα_o_ protein has impaired inhibitory signaling in cerebellar PCs, which is most likely due to increased levels of free Gβγ in inhibitory neurons.

Movement disorders may originate from abnormalities in several brain regions including striatum, cortex, and cerebellum. Cerebellar PCs have abundant G_o_ expression (Asano, Semba, Kamiya, Ogasawara, & Kato, 1988) so cerebellar PCs were the main focus of this study. Mutant mice exhibited decreased inhibitory signaling to PCs. The frequencies of sIPSCs and especially mIPSCs were reduced without changes in amplitude suggesting a pre-synaptic mechanism.

GABA_B_ receptor-activated Gα_o_ is the primary mediator of inhibition of the non-AP related-mIPSCs (Harvey & Stephens, 2004), however, the GABA_B_ receptor does not appear to be necessary for the actions of the mutant-dependent presynaptic inhibition. The mutant did not cause a leftward shift in the concentration-response curve for baclofen on the mIPSC frequency. More tellingly, the GABA_B_R antagonist CGP36214 did not alter the baseline mIPSC frequency in either WT or *Gnao1* mice at concentrations that prevented baclofen-induced inhibition of mIPSC frequency.

Moreover, the level of mutant Gα_o_ is about half that of WT in multiple brain regions including cerebellum. All these results are consistent with a haploinsufficient LOF *Gnao1* mutant function. With low levels of Gα_o_ (the most abundant Gα subunit (Jiang & Bajpayee, 2009)) and normal amounts of Gβγ, one would expect increased free Gβγ due to altered subunit stoichiometry. The enhanced free Gβγ could directly inhibit synaptic vesicle release through actions on SNARE complex or Ca^2+^ influx (Feng et al., 2018; Zurawski et al., 2017; Zurawski, Rodriguez, Hyde, Alford, & Hamm, 2016). The lack of effect of calcium channel block by Cd^2+^ on the mIPSC regulation would be consistent with this mechanism. One potentially incongruous finding in this study is that Gα_o_ blockers, PTX, significantly increased the mIPSC frequency in *Gnao1* mutant mice, while one would expect the opposite for a LOF *Gnao1* mutant. Although PTX is selective for Gα_o/i_, it can also lead to changes that may ultimately increase GABA release. For example, PTX locks Gα_o_ into an inactive state which makes it unable to release Gβγ to inhibit adenylyl cyclase (AC), causing a cAMP accumulation (Mangmool & Kurose, 2011). Rises in cAMP activate PKA, which enhances release probability at many synapses (Leenders & Sheng, 2005). Alternatively, PTX-mediated ADP-ribosylation of Gα_o_/βγ could stabilize the GDP-bound heterotrimer and reduce free Gβγ resulting from spontaneous GTP binding to Gα_o_ or could recruit Gβγ to Gα_i_ subunits (Figure 10).

**Figure 10.**
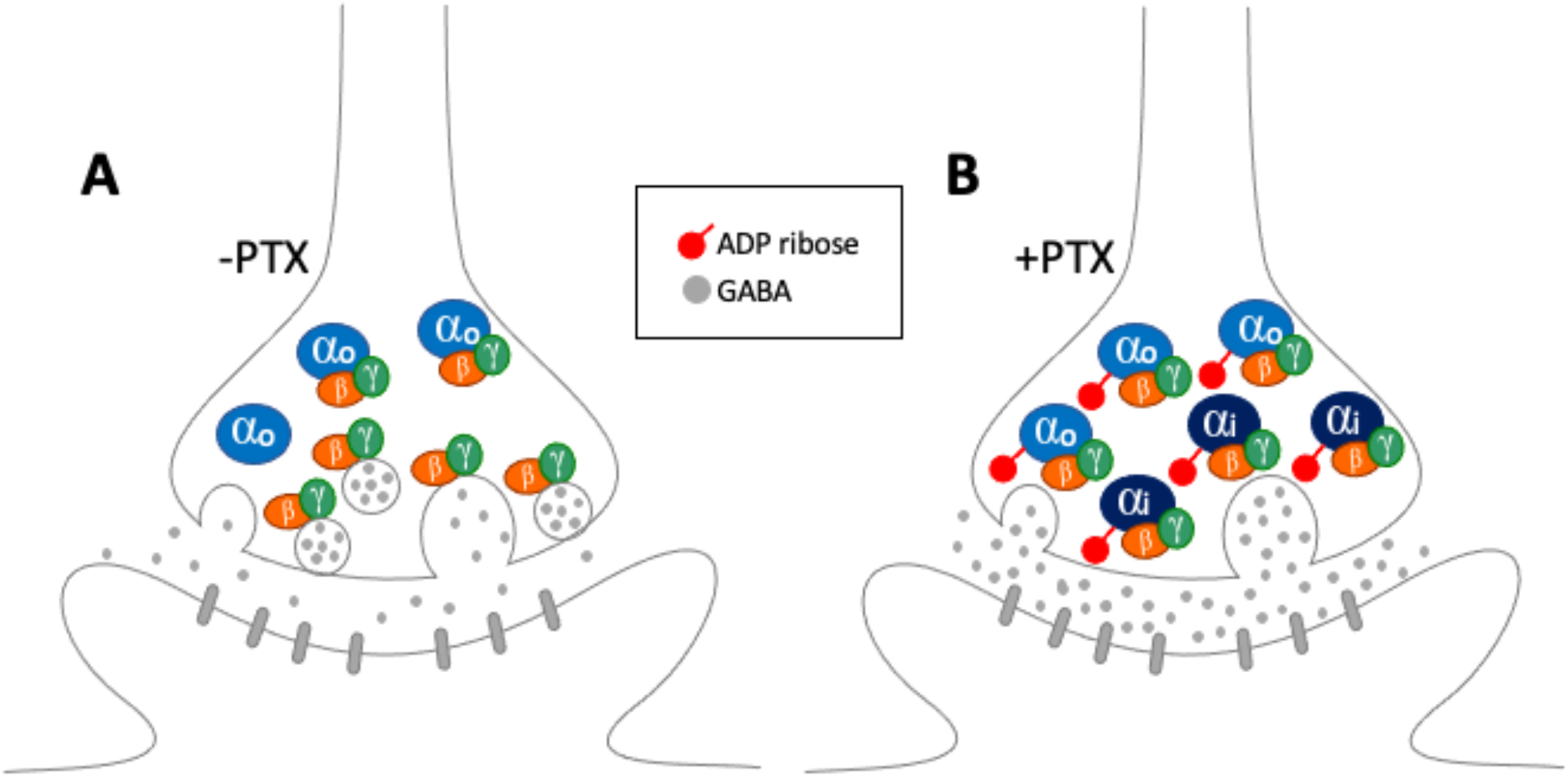
Model of Gα_o_ loss-of-function in suppression of inhibitory neurotransmitter release.

The limited role for receptors in the neurophysiological alterations in cerebellum reported here provide a cautionary tale for the suggestion that we have made for using GPCR antagonists to ameliorate movement disorders in patients (Feng et al. 2017; Feng et al. 2018). It is possible, however, that different signal outputs or brain regions may be affected differently, and distinct mutant alleles may also show different patterns. Future studies exploring mutant *GNAO1* mechanisms in different brain regions and cell-types as well as exploring the multiple Gα_o_-mediated signal outputs with different mutant alleles will be needed to test the generality of the model proposed here.

## Supporting information

All supplemental figures

## Notes

Authors report no conflict of interest.

Funding sources: Support for this project was from The Bow Foundation and the MSU Clinical Translational Science Institute; support for HF was from the American Epilepsy Society; Support for AJR from MSU ASPET SURF

### Competing Interest Statement

The authors have declared no competing interest.

## References

Amat, S. B., Rowan, M. J. M., Gaffield, M. A., Bonnan, A., Kikuchi, C., Taniguchi, H., & Christie, J. M. (2017). Using c-kit to genetically target cerebellar molecular layer interneurons in adult mice. PLoS One, 12(6), e0179347. doi:10.1371/journal.pone.0179347

Ananth, A. L., Robichaux-Viehoever, A., Kim, Y. M., Hanson-Kahn, A., Cox, R., Enns, G. M., … Bernstein, J. A. (2016). Clinical Course of Six Children With GNAO1 Mutations Causing a Severe and Distinctive Movement Disorder. Pediatr Neurol, 59, 81–84. doi:10.1016/j.pediatrneurol.2016.02.018

Arya, R., Spaeth, C., Gilbert, D. L., Leach, J. L., & Holland, K. D. (2017). GNAO1-associated epileptic encephalopathy and movement disorders: c.607G>A variant represents a probable mutation hotspot with a distinct phenotype. Epileptic Disord, 19(1), 67–75. doi:10.1684/epd.2017.0888

Asano, T., Semba, R., Kamiya, N., Ogasawara, N., & Kato, K. (1988). Go, a GTP-binding protein: immunochemical and immunohistochemical localization in the rat. J Neurochem, 50(4), 1164–1169. doi:10.1111/j.1471-4159.1988.tb10588.x

Bardo, S., Robertson, B., & Stephens, G. J. (2002). Presynaptic internal Ca2+ stores contribute to inhibitory neurotransmitter release onto mouse cerebellar Purkinje cells. Br J Pharmacol, 137(4), 529–537. doi:10.1038/sj.bjp.0704901

Blackmer, T., Larsen, E. C., Takahashi, M., Martin, T. F., Alford, S., & Hamm, H. E. (2001). G protein betagamma subunit-mediated presynaptic inhibition: regulation of exocytotic fusion downstream of Ca2+ entry. Science, 292(5515), 293–297. doi:10.1126/science.1058803

Bologna, M., & Berardelli, A. (2018). The cerebellum and dystonia. Handb Clin Neurol, 155, 259–272. doi:10.1016/B978-0-444-64189-2.00017-2

Boxall, A. R. (2000). GABAergic mIPSCs in rat cerebellar Purkinje cells are modulated by TrkB and mGluR1-mediated stimulation of Src. J Physiol, 524 Pt 3, 677–684. doi:10.1111/j.1469-7793.2000.00677.x

Brown, A. M., Arancillo, M., Lin, T., Catt, D. R., Zhou, J., Lackey, E. P., … Sillitoe, R. V. (2019). Molecular layer interneurons shape the spike activity of cerebellar Purkinje cells. Sci Rep, 9(1), 1742. doi:10.1038/s41598-018-38264-1

Buijsen, R. A. M., Toonen, L. J. A., Gardiner, S. L., & van Roon-Mom, W. M. C. (2019). Genetics, Mechanisms, and Therapeutic Progress in Polyglutamine Spinocerebellar Ataxias. Neurotherapeutics, 16(2), 263–286. doi:10.1007/s13311-018-00696-y

Eccles, J. C. (1967). Circuits in the cerebellar control of movement. Proc Natl Acad Sci U S A, 58(1), 336–343. doi:10.1073/pnas.58.1.336

Feng, H., Khalil, S., Neubig, R. R., & Sidiropoulos, C. (2018). A mechanistic review on GNAO1-associated movement disorder. Neurobiol Dis. doi:10.1016/j.nbd.2018.05.005

Feng, H., Larrivee, C. L., Demireva, E. Y., Xie, H., Leipprandt, J. R., & Neubig, R. R. (2019). Mouse models of GNAO1-associated movement disorder: Allele-and sex-specific differences in phenotypes. PLoS One, 14(1), e0211066. doi:10.1371/journal.pone.0211066

Feng, H., Sjogren, B., Karaj, B., Shaw, V., Gezer, A., & Neubig, R. R. (2017). Movement disorder in GNAO1 encephalopathy associated with gain-of-function mutations. Neurology, 89(8), 762–770. doi:10.1212/WNL.0000000000004262

Fremont, R., Tewari, A., Angueyra, C., & Khodakhah, K. (2017). A role for cerebellum in the hereditary dystonia DYT1. Elife, 6. doi:10.7554/eLife.22775

Harvey, V. L., & Stephens, G. J. (2004). Mechanism of GABA receptor-mediated inhibition of spontaneous GABA release onto cerebellar Purkinje cells. Eur J Neurosci, 20(3), 684–700. doi:10.1111/j.1460-9568.2004.03505.x

Heintz, N. (2004). Gene expression nervous system atlas (GENSAT). Nat Neurosci, 7(5), 483. doi:10.1038/nn0504-483

Hirota, S., Ito, A., Morii, E., Wanaka, A., Tohyama, M., Kitamura, Y., & Nomura, S. (1992). Localization of mRNA for c-kit receptor and its ligand in the brain of adult rats: an analysis using in situ hybridization histochemistry. Brain Res Mol Brain Res, 15(1-2), 47–54. doi:10.1016/0169-328x(92)90150-a

Ikeda, S. R. (1996). Voltage-dependent modulation of N-type calcium channels by G-protein beta gamma subunits. Nature, 380(6571), 255–258. doi:10.1038/380255a0

Jiang, M., & Bajpayee, N. S. (2009). Molecular mechanisms of go signaling. Neurosignals, 17(1), 23–41. doi:10.1159/000186688

Kelly, M., Park, M., Mihalek, I., Rochtus, A., Gramm, M., Perez-Palma, E., … Poduri, A. (2019). Spectrum of neurodevelopmental disease associated with the GNAO1 guanosine triphosphate-binding region. Epilepsia, 60(3), 406–418. doi:10.1111/epi.14653

Keshet, E., Lyman, S. D., Williams, D. E., Anderson, D. M., Jenkins, N. A., Copeland, N. G., & Parada, L. F. (1991). Embryonic RNA expression patterns of the c-kit receptor and its cognate ligand suggest multiple functional roles in mouse development. EMBO J, 10(9), 2425–2435. Retrieved from https://www.ncbi.nlm.nih.gov/pubmed/1714375

Kirshner, H., Aguet, F., Sage, D., & Unser, M. (2013). 3-D PSF fitting for fluorescence microscopy: implementation and localization application. J Microsc, 249(1), 13–25. doi:10.1111/j.1365-2818.2012.03675.x

Larrivee, C. L., Feng, H., Quinn, J. A., Shaw, V. S., Leipprandt, J. R., demireva, E. Y., … Neubig, R. R. (2019). Preprint: Mice with GNAO1 R209H Movement Disorder Variant Display Hyperlocomotion Alleviated by Risperidone. BioRxiv. Retrieved from https://doi.org/10.1101/662031

Leenders, A. G., & Sheng, Z. H. (2005). Modulation of neurotransmitter release by the second messenger-activated protein kinases: implications for presynaptic plasticity. Pharmacol Ther, 105(1), 69–84. doi:10.1016/j.pharmthera.2004.10.012

Mangmool, S., & Kurose, H. (2011). G(i/o) protein-dependent and -independent actions of Pertussis Toxin (PTX). Toxins (Basel), 3(7), 884–899. doi:10.3390/toxins3070884

Manova, K., Bachvarova, R. F., Huang, E. J., Sanchez, S., Pronovost, S. M., Velazquez, E., … Besmer, P. (1992). c-kit receptor and ligand expression in postnatal development of the mouse cerebellum suggests a function for c-kit in inhibitory interneurons. J Neurosci, 12(12), 4663–4676. Retrieved from https://www.ncbi.nlm.nih.gov/pubmed/1281492

Motro, B., van der Kooy, D., Rossant, J., Reith, A., & Bernstein, A. (1991). Contiguous patterns of c-kit and steel expression: analysis of mutations at the W and Sl loci. Development, 113(4), 1207–1221. Retrieved from https://www.ncbi.nlm.nih.gov/pubmed/1811937

Nakamura, K., Kodera, H., Akita, T., Shiina, M., Kato, M., Hoshino, H., … Saitsu, H. (2013). De Novo mutations in GNAO1, encoding a Galphao subunit of heterotrimeric G proteins, cause epileptic encephalopathy. Am J Hum Genet, 93(3), 496–505. doi:10.1016/j.ajhg.2013.07.014

Reeber, S. L., Otis, T. S., & Sillitoe, R. V. (2013). New roles for the cerebellum in health and disease. Front Syst Neurosci, 7, 83. doi:10.3389/fnsys.2013.00083

Saitsu, H., Fukai, R., Ben-Zeev, B., Sakai, Y., Mimaki, M., Okamoto, N., … Matsumoto, N. (2016). Phenotypic spectrum of GNAO1 variants: epileptic encephalopathy to involuntary movements with severe developmental delay. Eur J Hum Genet, 24(1), 129–134. doi:10.1038/ejhg.2015.92

Schindelin, J., Arganda-Carreras, I., Frise, E., Kaynig, V., Longair, M., Pietzsch, T., … Cardona, A. (2012). Fiji: an open-source platform for biological-image analysis. Nat Methods, 9(7), 676–682. doi:10.1038/nmeth.2019

Schorling, D. C., Dietel, T., Evers, C., Hinderhofer, K., Korinthenberg, R., Ezzo, D., … Kirschner, J. (2017). Expanding Phenotype of De Novo Mutations in GNAO1: Four New Cases and Review of Literature. Neuropediatrics, 48(5), 371–377. doi:10.1055/s-0037-1603977

Sunahara, R. K., & Taussig, R. (2002). Isoforms of mammalian adenylyl cyclase: multiplicities of signaling. Mol Interv, 2(3), 168–184. doi:10.1124/mi.2.3.168

Vanni, V., Puglisi, F., Bonsi, P., Ponterio, G., Maltese, M., Pisani, A., & Mandolesi, G. (2015). Cerebellar synaptogenesis is compromised in mouse models of DYT1 dystonia. Exp Neurol, 271, 457–467. doi:10.1016/j.expneurol.2015.07.005

Walker, R. H. (2016). The non-Huntington disease choreas: Five new things. Neurol Clin Pract, 6(2), 150–156. doi:10.1212/CPJ.0000000000000236

Yuan, Y., & Atchison, W. D. (1999). Comparative effects of methylmercury on parallel-fiber and climbing-fiber responses of rat cerebellar slices. J Pharmacol Exp Ther, 288(3), 1015–1025. Retrieved from https://www.ncbi.nlm.nih.gov/pubmed/10027838

Yuan, Y., & Atchison, W. D. (2003a). Electrophysiological studies of neurotoxicants on central synaptic transmission in acutely isolated brain slices. Curr Protoc Toxicol, Chapter 11, Unit11 11. doi:10.1002/0471140856.tx1111s17

Yuan, Y., & Atchison, W. D. (2003b). Methylmercury differentially affects GABA(A) receptor-mediated spontaneous IPSCs in Purkinje and granule cells of rat cerebellar slices. J Physiol, 550(Pt 1), 191–204. doi:10.1113/jphysiol.2003.040543

Yuan, Y., & Atchison, W. D. (2007). Methylmercury-induced increase of intracellular Ca2+ increases spontaneous synaptic current frequency in rat cerebellar slices. Mol Pharmacol, 71(4), 1109–1121. doi:10.1124/mol.106.031286

Yuan, Y., & Atchison, W. D. (2016). Multiple Sources of Ca2+ Contribute to Methylmercury-Induced Increased Frequency of Spontaneous Inhibitory Synaptic Responses in Cerebellar Slices of Rat. Toxicol Sci, 150(1), 117–130. doi:10.1093/toxsci/kfv314

Zhang, Q., Dickson, A., & Doupnik, C. A. (2004). Gbetagamma-activated inwardly rectifying K(+) (GIRK) channel activation kinetics via Galphai and Galphao-coupled receptors are determined by Galpha-specific interdomain interactions that affect GDP release rates. J Biol Chem, 279(28), 29787–29796. doi:10.1074/jbc.M403359200

Zurawski, Z., Page, B., Chicka, M. C., Brindley, R. L., Wells, C. A., Preininger, A. M., … Hamm, H. E. (2017). Gbetagamma directly modulates vesicle fusion by competing with synaptotagmin for binding to neuronal SNARE proteins embedded in membranes. J Biol Chem, 292(29), 12165–12177. doi:10.1074/jbc.M116.773523

Zurawski, Z., Rodriguez, S., Hyde, K., Alford, S., & Hamm, H. E. (2016). Gbetagamma Binds to the Extreme C Terminus of SNAP25 to Mediate the Action of Gi/o-Coupled G Protein-Coupled Receptors. Mol Pharmacol, 89(1), 75–83. doi:10.1124/mol.115.101600

