## Supplementary material for "Mice With Monoallelic *GNAO1* Loss Exhibit Reduced Inhibitory Synaptic Input To Cerebellar Purkinje Cells": All supplemental figures

### EXTENDED DATA

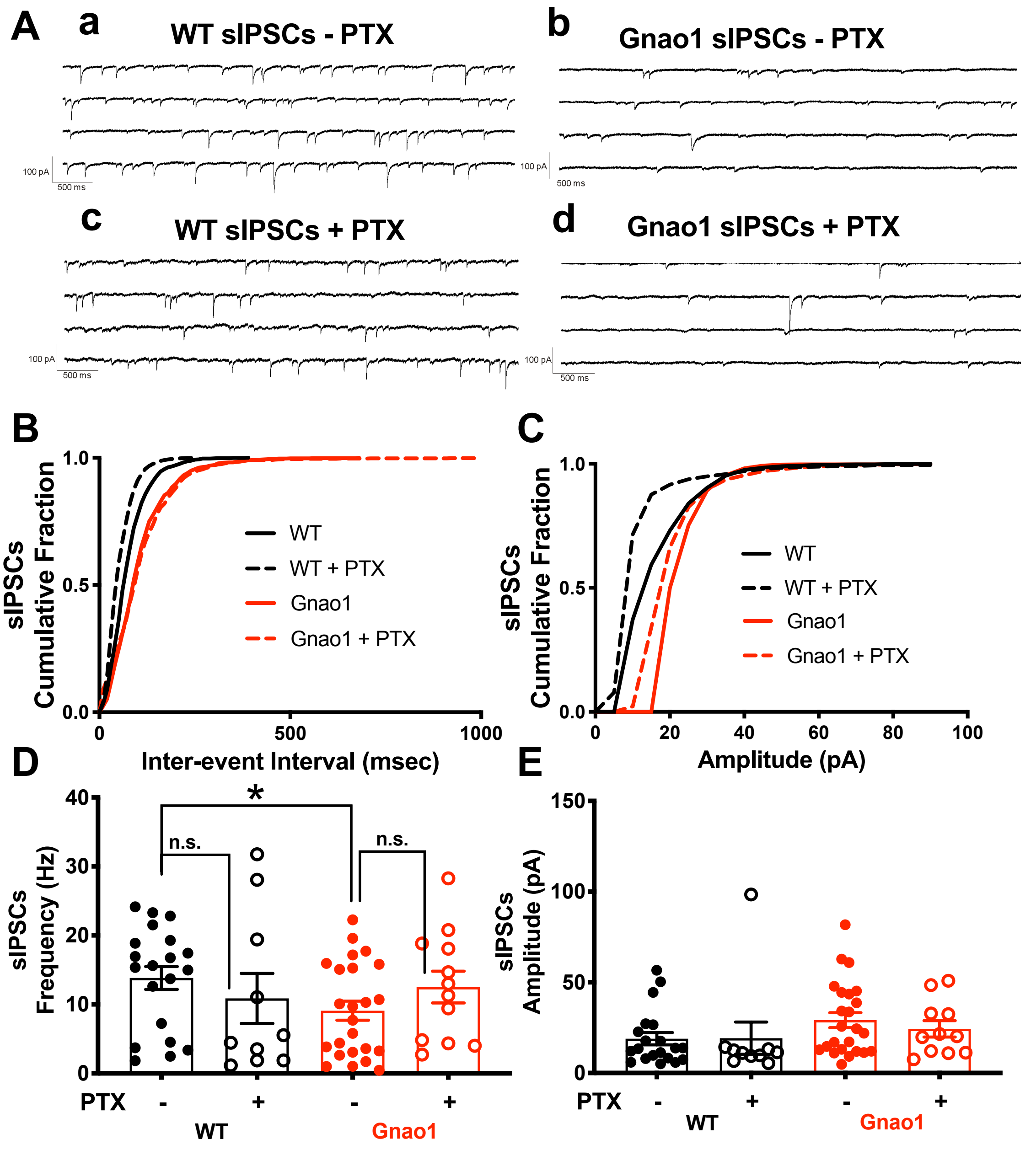


**Figure 5-1. PTX did not affect either frequency or amplitude of sIPSCs in WT and Gnao1 mice.** Slices were incubated in 1 μM/ml of PTX for >6 hrs pre-recording. Representative traces showed the example recordings of (A) sIPSCs in WT and Gnao1 mice before and after PTX incubation. Neither frequency (B, D) nor amplitude (C, E) of sIPSCs was affected by PTX incubation. Unpaired Student's t-test; WT (n=5 mice), Gnao1 (n=6 mice); *p=0.03 between WT vs. Gnao1 mice without PTX incubation. Results between WT and Gnao1 were not significant.

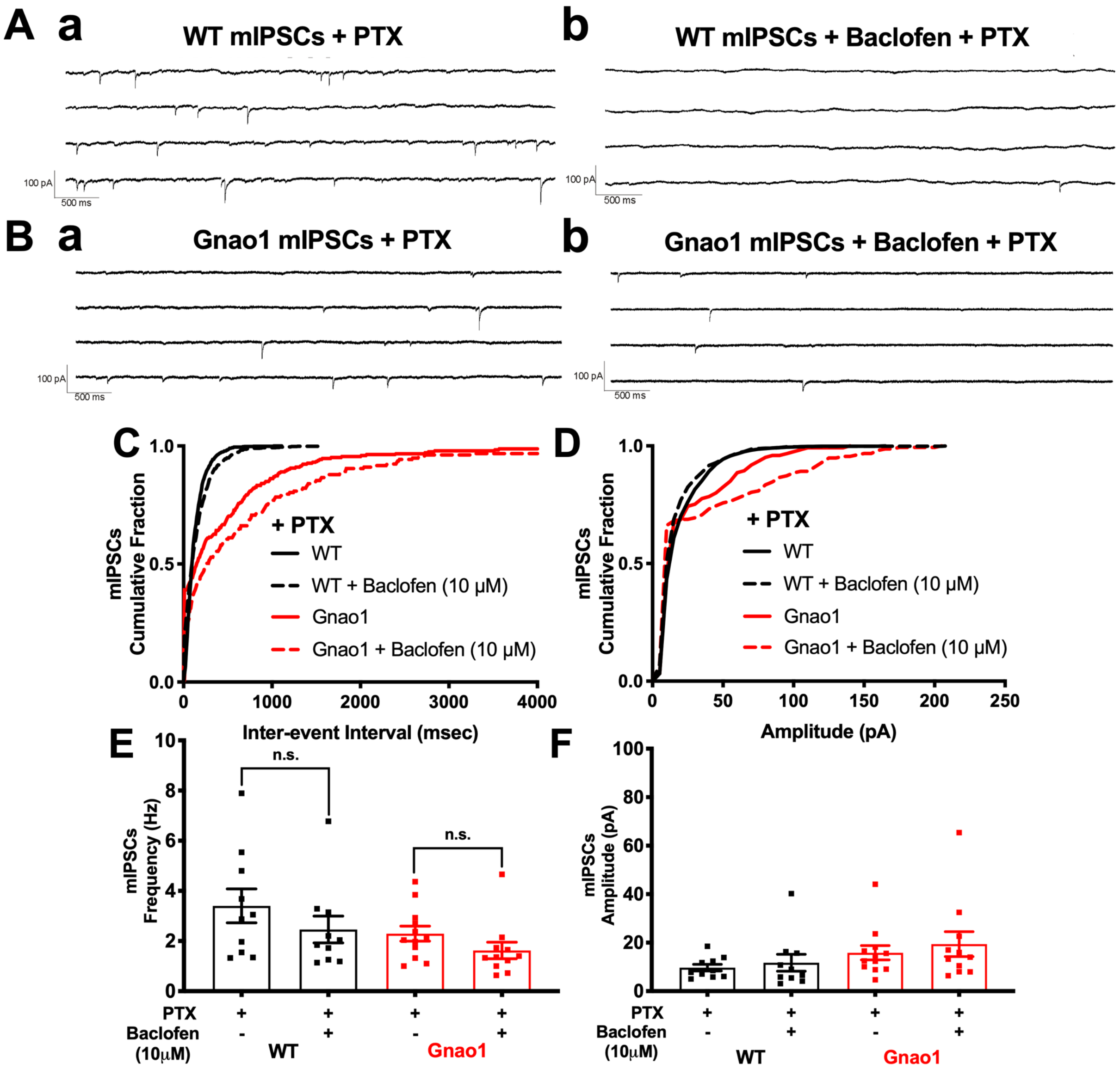


**Figure 6-1. PTX incubation eliminates baclofen-induced inhibition of mIPSC frequency in WT and Gnao1 mutant mice.** No significant change was observed in the (C, E) frequency or the (D, F) amplitude of mIPSCs after adding baclofen in both WT and Gnao1 mice with PTX incubation. Unpaired Student's t-test; WT (n=5), Gnao1 (n=6).

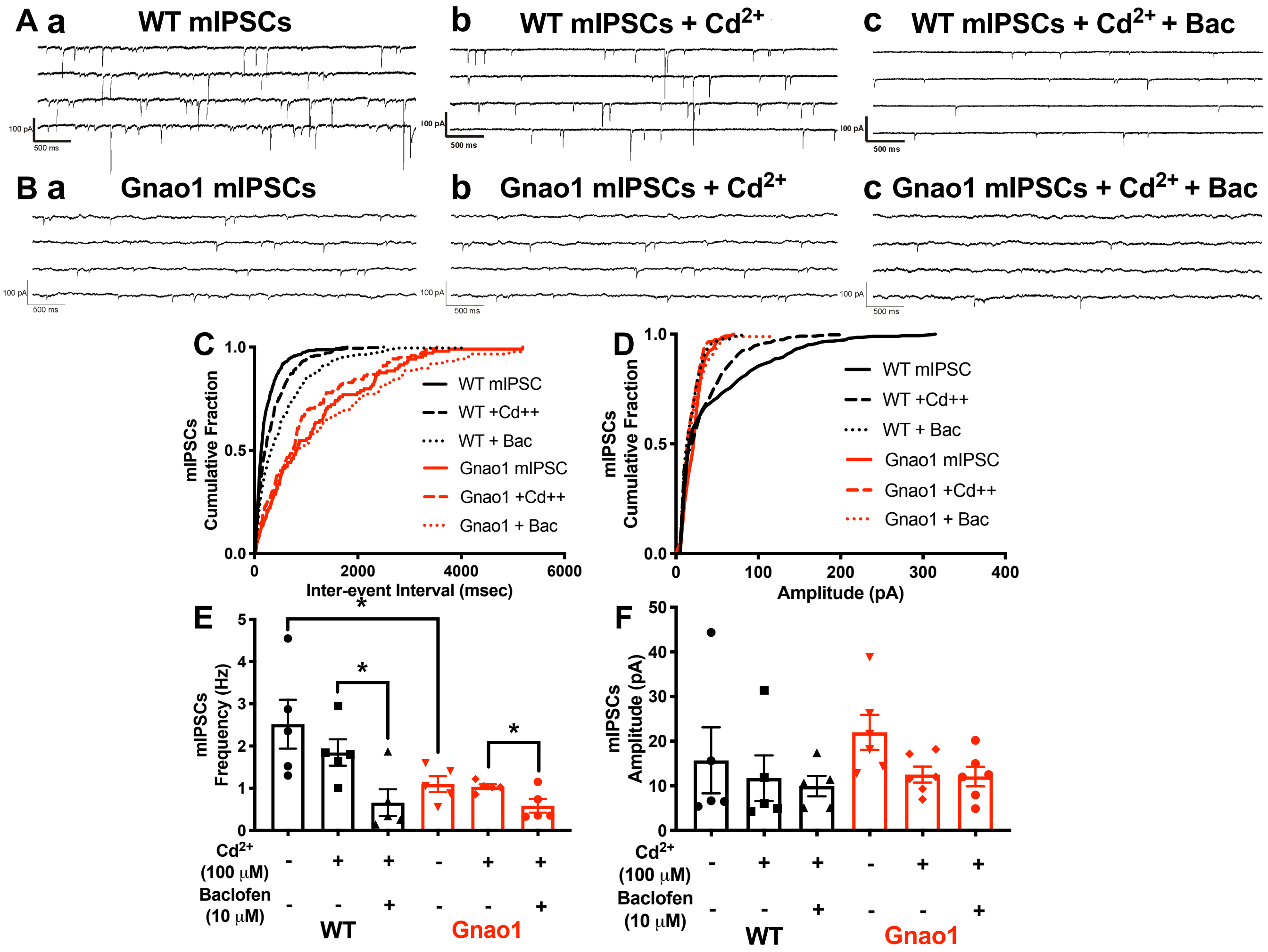


**Figure 6-2. Cadmium-block of extracellular calcium influx does not significantly affect the frequency or the amplitude of mIPSCs in either WT and Gnao1 mutant mice.** Representative recordings showing the sIPSCs of (A) WT and (B) Gnao1 in (a) ASCF with 100 µM AP-V, 10 µM CNQX and 0.5 µM TTX; (b) $Cd^{2+}$(100 µM)-ASCF with AP-V, CNQX and TTX; (c) $Cd^{2+}$ (100 µM)-ASCF with AP-V, CNQX, TTX and 10 µM baclofen. (C, E) 100 µM $Cd^{2+}$ did not reduce the frequency of mIPSCs in either WT or Gnao1 mice. Baclofen (10 µM) reduced the frequency of mIPSC with the presence of 100 µM $Cd^{2+}$. (D, F) Similarly, 100 µM $Cd^{2+}$ does not affect amplitudes of sIPSCs. Unpaired Student's t-test; WT (n=5 mice), Gnao1 (n=5 mice); $^*p<0.05$.

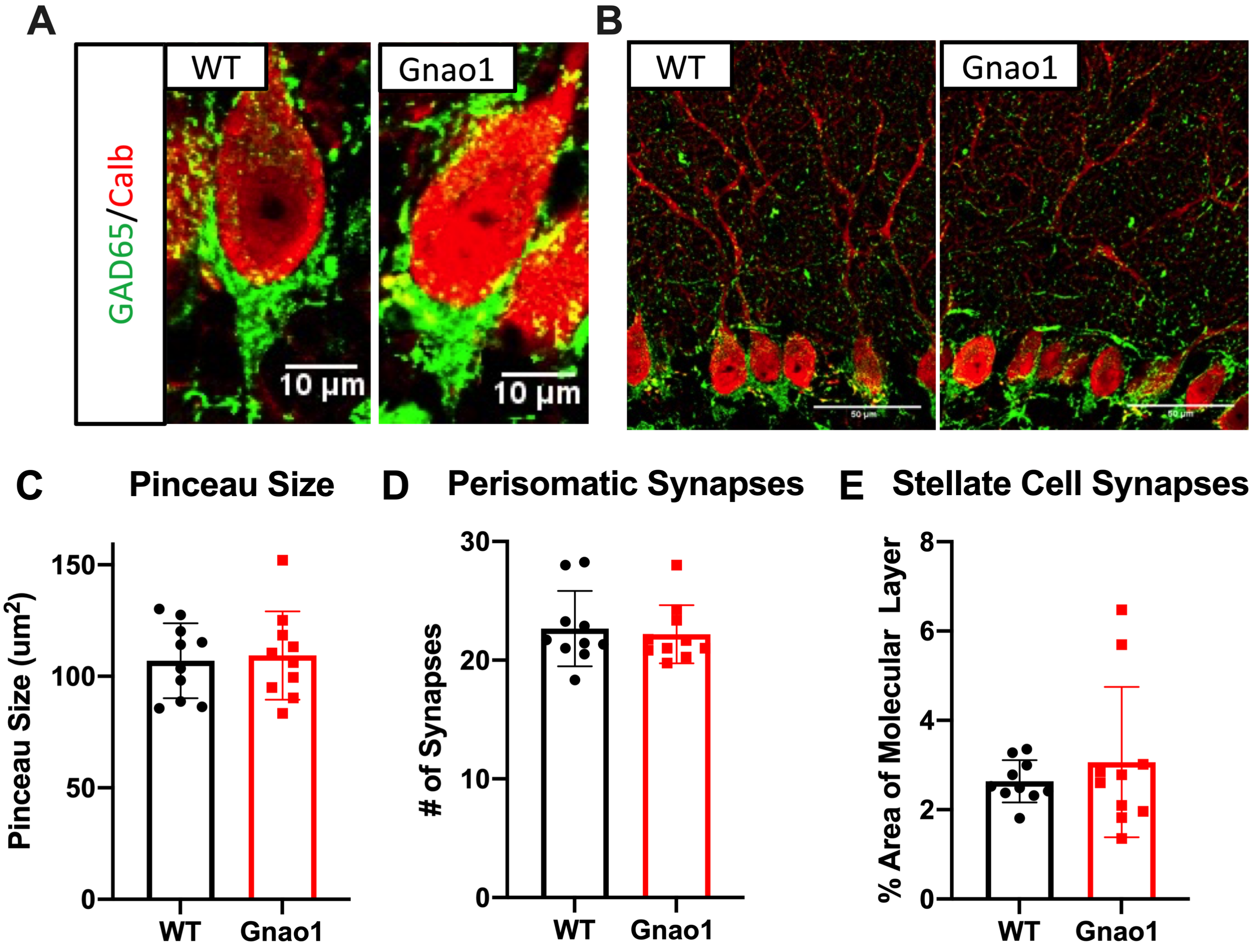


**Figure 9-1. Basket- and stellate-Purkinje synapses show no difference in *Gnao1* mutants.** (A) Representative images of GAD65 staining at WT and *Gnao1* mutant basket cell-PC synapses. (B) Representative views of WT and *Gnao1* mutant molecular layers, with GAD65 staining at stellate cell-PC synapses. (C,D) No significant difference was observed in the pinceau size (n=147 cells) or number of perisomatic puncta (n=128 cells) between WT and *Gnao1* mutant mice; WT (n=10), Gnao1 (n=10). Measurements were taken at a PC's widest point. (E) There was no significant difference in the percent of the molecular layer occupied by GAD65 puncta. Some off-target vasculature staining is visible, which was corrected for by excluding particles larger than 3 $\mu m^2$.

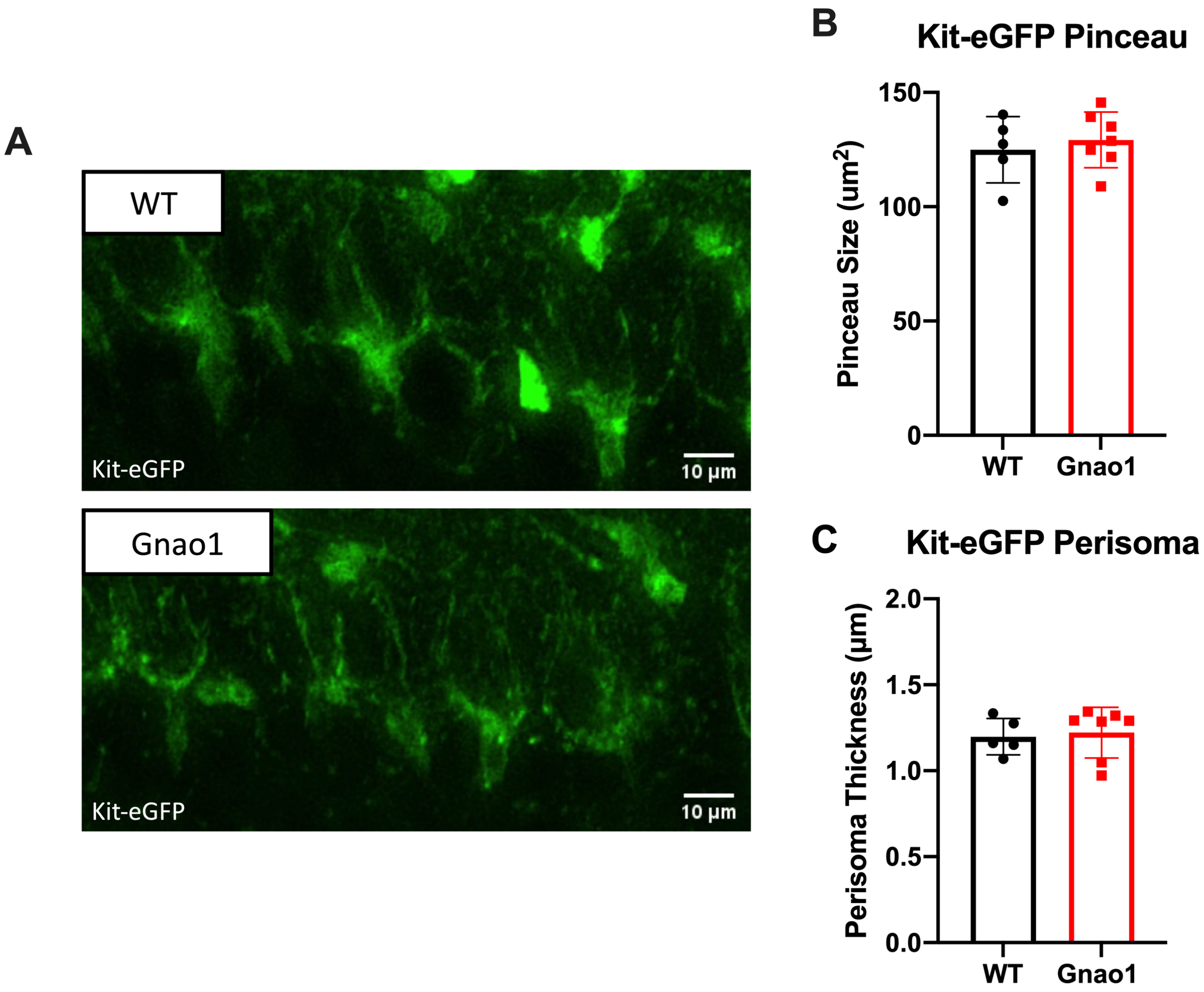


**Figure 9-2. Basket axon contacts with PCs are normal in *Gnao1* mutants.** (A) Representative images of the PC layer in WT Kit-eGFP and *Gnao1* mutant x Kit-eGFP mice. Molecular layer interneuron axon collaterals are marked by Kit-eGFP. (B,C) No differences in pinceau size (n=353 cells) or perisomatic contacts (n=453 cells) were observed between WT and *Gnao1* mutant mice; WT (n=5), Gnao1 (n=7).
